# Failure of DNA repair leads to the tumorigenic transformation of a regenerating blastema in *Drosophila* imaginal discs

**DOI:** 10.1101/2025.10.24.684460

**Authors:** Tanroop Kaur, Ada Repiso, Ying Liu, Paula Doria, Joan Urgell, Luis Rodriguez-Escudero, Katerina Karkali, Yanhui Hu, Norbert Perrimon, Enrique Martin-Blanco

## Abstract

We present a *Drosophila* model demonstrating that repeated cycles of tissue regeneration following localized apoptosis can induce tumorigenesis in the absence of oncogene activation or tumor suppressor loss. Chronic regenerative stress leads to epithelial disorganization, loss of cell polarity, immune cell infiltration, DNA damage, centrosomal defects, and aneuploidy— classical hallmarks of cancer. A subset of “walking dead” cells resists apoptosis and contributes to tissue overgrowth and heterogeneity. Single-cell transcriptomics identifies a distinct cell population, absent from normal tissue, emerging under both regenerative and tumorigenic conditions and characterized by enrichment in JNK pathway components. The tumorigenic signature specifically features distinctive JNK regulatory elements while excluding chaperone proteins. Notably, inhibition of JNK signaling or modulation of DNA repair suppresses neoplastic transformation. This model reveals how chronic injury and defective regeneration drive cancer initiation, establishing a mechanistic link between wound healing and tumorigenesis, and providing a powerful system to explore early tumor development and therapeutic intervention.

## INTRODUCTION

There is mounting evidence that many tumors arise from the transformation of normal stem cells into cancer cells with a virtually infinite capacity for self-renewal ^1^. This tumor-propagating potential may relate to the cancer stem cells ability to hijack natural self-renewal programs, such as those activated during tissue regeneration ^2^.

Successful regeneration, in common to natural morphogenetic events, requires a tightly regulated control of cell proliferation, polarity, and patterning ^3^. If one or more of these processes become uncontrolled, unsuccessful regeneration may result in tumorigenesis. Indeed, tumors have been described as “wounds that won’t heal,” reflecting the striking similarities between cancerous tumor growth and wound healing ^4^. Numerous cases of wound-related cancers have been reported, and a close association between chronic tissue damage, failed regeneration, inflammation, and cancer has been repeatedly observed ^5^. Despite these well documented connections, it is still lacking a detailed survey of the communalities and differences between tissue repair and tumor formation under stress.

Gene conservation allows model organisms to be used in studying genetic pathologies, and this is instrumental when comparing wound healing or tumorigenesis. *Drosophila* imaginal discs, which are in many aspects comparable to mammalian epithelia, are constituted by a monolayer epithelium with apical-basal polarity limited by a basal extracellular matrix. Importantly, they serve as an excellent model for tissue repair studies as both, mechanical injuries and genetic ablations lead disc cells to change fate and to implement a complete regeneration program. Cultured disc fragments following implantation into the abdomen of adult females also regenerate ^6^. In the wing disc, multiple studies have focused on the exploration of the molecular mechanisms and cellular processes operational on the early aspects of healing and on the late regenerative responses ^7–9^. Evolutionarily conserved genes essential for healing have been identified ^7,10^ and strong parallels between the early stages of regeneration and tumor formation have been observed ^11^. Yet, key aspects such as regeneration’s fidelity remain unexplored. Regeneration is highly accurate and the mechanisms ensuring precision have not been evaluated. We hypothesized that by manipulating regenerating cells we could study the mechanisms controlling this precision and identify the factors that, when these mechanisms are challenged, trigger regeneration failure or, eventually, tumor formation. In this context, the *Drosophila* imaginal discs also constitute a reliable system to model the onset and progression of epithelial cancers. The activation of the Notch pathway or the overexpression of the oncogenic form of Ras, RasV12, in these discs result in epithelial hyperplasia. This type of benign tumors can be driven into malignancy in different mutant backgrounds affecting tumor suppressors like *scrib*, *Dlg* or *Lgl*, which affect cell polarity. These malignant tumors show tissue overgrowth, loss of apicobasal polarity and invasive and metastatic capabilities ^12–14^. *Drosophila* tumors in the wing discs have also been observed after the overexpression of the epidermal growth factor receptor (EGFR) or in mutants of the Hippo tumor-suppressor pathway. In this last case associated to elevated apoptosis ^15^. Remarkably, mutations on all these genes are involved in the development of malignant tumors in mammals.

In this report, we demonstrate that inducing multiple regeneration events through repeated localized apoptosis induction leads to uncontrolled cell proliferation. The overproliferating tissue loses apicobasal polarity and planar patterning and is infiltrated by migratory hemocytes through a disrupted basal lamina. These discs exhibit high levels of DNA double-strand breaks (DSBs), widespread centrosomal aberrations, and aneuploidies. The neoplastic overgrowths observed after repetitive local cell death induction comprise heterogeneous cell populations, including cells that, challenging apoptosis, behave as “walking dead cells,” intercalating throughout the tissue to distant positions. Moreover, non-cell-autonomous effects are observed in wild-type cells that are induced to proliferate. Single cell transcriptomic analysis identifies a novel cell subpopulation absent in wild-type unstressed discs in both, regenerative and tumorigenic conditions. This subpopulation (SPIKE) in neoplastic discs is distinctly enriched in genes involved in or regulating the JNK signaling cascade. Remarkably, the neoplastic transformations resulting from repeated cell death insults are rescued by interfering with JNK activity or manipulating DNA repair responses.

In summary, we have developed a model of tumorigenesis independent of oncogene activation or tumor suppressor inactivation. This model offers a powerful tool for understanding tumor development in tissues subjected to chronic stress.

## RESULTS

### Induction of uncontrolled growth in regenerative tissues by repeated stimulation of cell death

Temporally restricted Gal4-mediated induction of cell death in *Drosophila* third instar larvae (by overexpressing the pro-apoptotic gene *reaper* (*rpr*)), results in complete tissue regeneration^16^. If Ptc-Gal4-driven cell death is induced for 16 h only (in a stripe of cells at the center of the wing disc (96 h of development)) employing the temperature sensitive repressor of Gal4, Gal80^TS^, the animals manage to regenerate the affected tissue and develop a normal wing [Regeneration Induction 1 (RI1). Taking advantage of this well-established protocol, we asked if interfering in the regeneration process, by the sequential and repetitive induction of apoptosis, could alter its fidelity and efficiency. Remarkably, three (RI3) or more rounds of timely controlled death-induction and recovery (**Figure 1A**) lead to the reprogramming of the regenerative tissue to an over proliferative uncontrolled regime. Wing discs show, with a penetrance of 10-20%, a distinguishable unpatterned overgrowth by the end of the larval period, not limited to the location of the challenged tissue, resulting in the formation of ectopic imaginal epithelia folds (compare **Figures 1B** and **1C**). Adults never eclose and pharates show different types of aberrations (not shown).

**Figure 1.**
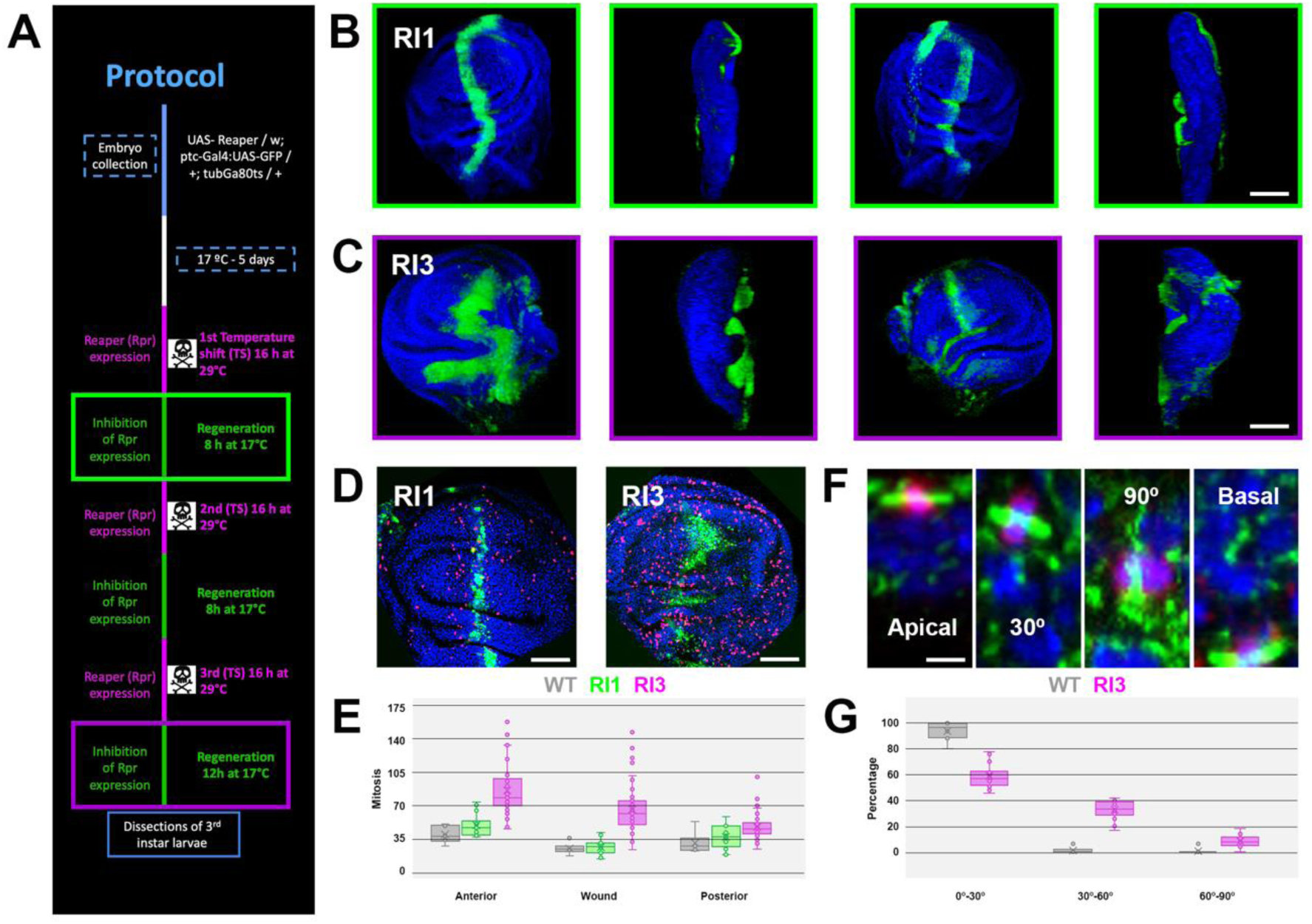
Sequential apoptosis induction leads to uncontrolled cell proliferation of a regenerating blastema. **A)** Experimental protocol of tumorigenic induction. Temperature controlled (tubGal80^ts^) repetitive induction of Rpr expression along the antero/posterior compartment border (Ptc expression domain) in wing imaginal discs leads in 3 series to uncontrolled tissue overgrowths. Text in magenta highlights apoptosis induction events. Text in green points to death recovery periods. Green box indicates initial successful regeneration (RI1) (wing disc in **B**). Magenta box denotes aberrant tumorigenic transformation (RI3) (wing disc in **C**). **B)** Frontal lateral and back views of a regenerating wing disc (RI1). Green is GFP under the control of Ptc and blue shows nuclear DAPI staining. Scale bar is 50 µm. GFP expression and wing disc and shape are identical to those of wild type discs not expressing Rpr (not shown). **C)** Frontal lateral and back views of a tumorigenic wing disc (RI3). Color code and scale bar as in B. **D)** H3^P^ labelling (magenta) of RI1 and RI3 wing imaginal disc depicting cells in mitosis. Green is GFP and blue is DAPI. Scale bar is 40 µm. **E)** Quantification of mitotic cells in wild type (grey) (n = 6), RI1 (green) (n = 15) and RI3 (magenta) (n = 47) wing imaginal discs in the anterior, wound domain (Ptc/GFP) and posterior wing compartments. Two-tailed P values between WT and RI / Rpr are not significant in any domain (0,0741, 0,7104 and 0,2637). P values both between WT and RI3 / Rpr and between RI / Rpr and RI3 / Rpr (0,0002, 0,0003 and 0.0039; and <0.0001, <0.0001 and 0,0079 respectively) are all highly statistically significant. **F)** Mitotic orientation of cell divisions Individual examples from left to right of planar apical, tilted (30°), perpendicular (90°) and basal divisions. Green is alpha-tubulin, red is H3^P^ and blue is DAPI. Scale bar is 5 µm. **G)** Quantification of frequencies of mitotic orientation [in percentage in 3 categories (0°-30°, 30°-60° and 60°-90°)] for wild type (grey) (n = 8) and RI3 (magenta) ( n = 17) wing imaginal discs.

It is known that regenerating wing imaginal discs, after one round of Rpr 16 h induced apoptosis, show a peak of mitoses immediately after death induction ^17^. To test whether the observed overgrowths were caused by an increase in the number of cell divisions we analyzed the number of cells in mitosis employing an anti-PH3 antibody. We analyzed samples challenged only once and compared them to samples challenged three and four times, plus or minus a final 12 h recovery period. Upon single death induction in the Ptc domain, cell division numbers increase in both, those cells located in the Ptc domain and also those in the anterior compartment of the wing disc, while cells from the posterior compartment don’t respond ^16^. We found that the topological distribution of mitosis was conserved upon three death/regeneration episodes but the scalar drastically changed and a huge increase in mitosis number was observed. Cells’ divisions become unrestrained and they keep multiplying overtime with no control (**Figures 1D** and **1E**).

Mitotic division typically initiates with cell rounding, which in polarized epithelia, is generally followed by planar alignment of the mitotic spindle and cell cleavage orthogonal to the plane of the epithelium ^18–22^. The disruption of planar divisions has been proposed to cause epithelial to mesenchymal transition and cancer ^23–25^. Experimentally induced non-planar divisions correlate with cell delamination and apoptotic death, and blocking the death becomes sufficient to drive the formation of basally localized tumor-like masses ^22^. In *Drosophila* regenerating wing discs, cells’ divisions always orient planarly towards the wound. We found that multiple rounds of death/regeneration elicit, not just an increase in proliferation, but also the loss of planar alignment and, occasionally the mislocalization of planar divisions to basal positions (**Figure 1F** and **1G**).

We conclude that, when sequentially challenged multiple times to death and regeneration, the wing imaginal disc is unable to control mitosis rates and cell divisions orientation leading to the formation of hyperplastic overgrowths.

### Loss of apicobasal polarity of the regenerating blastema

Unregulated overgrowth by itself does not necessarily imply the malignant transformation of a tissue. Key aspects of cancer cells include evasion of apoptosis, self-sufficiency in growth signals, insensitivity to anti-growth signals, sustained angiogenesis, unlimited replicative potential, invasiveness and metastasis ^26^. Additionally, in epithelia, these landmarks include events such as the loss of apicobasal polarity ^27^ and the degradation of the Extra Cellular Matrix (ECM) ^28^.

In imaginal discs the epithelia monolayer displays stereotyped apical and basolateral membrane domains and is limited by a basal ECM. Three protein complexes establish and maintain discs’ epithelial polarity: the Crumbs/Stardust/PATJ/Bazooka, the Par6/aPKC and the Scrib/Dlg/Lgl complexes, which are respectively placed at the apical, subapical and basolateral membrane domains ^29^. importantly, all the members of this last one have been shown to act as neoplastic Tumor Suppressors ^30^.

We monitored apicobasal polarity in the regenerating blastema employing anti-Crumbs and anti-Dlg antibodies and found that the sequential induction of multiple rounds (three) of cell death resulted in the scattered loss of apicobasal components from the overgrowing imaginal tissue, which constitute a compelling proof of their neoplastic character. Crumbs is expressed lining the apical domain of all imaginal and peripodial epithelial cells of the wing disc. This expression fades away and multiple gaps were observed in response to sequential cell death rounds (compare **Figures 2A** to **2B**). Likewise, equivalent breaks in the expression of Dlg, which decorates the whole epithelia in the wild type, were also frequently observed (compare **Figures 2C** to **2D**). The impairment of apicobasal polarity in the neoplastic regenerating blastema correlates with a loss of epithelial integrity and E-cadherin, the most important component of Adherens Junctions (AJs) ^31^, was also downregulated (compare **Figure 2E** to **2F**).

**Figure 2.**
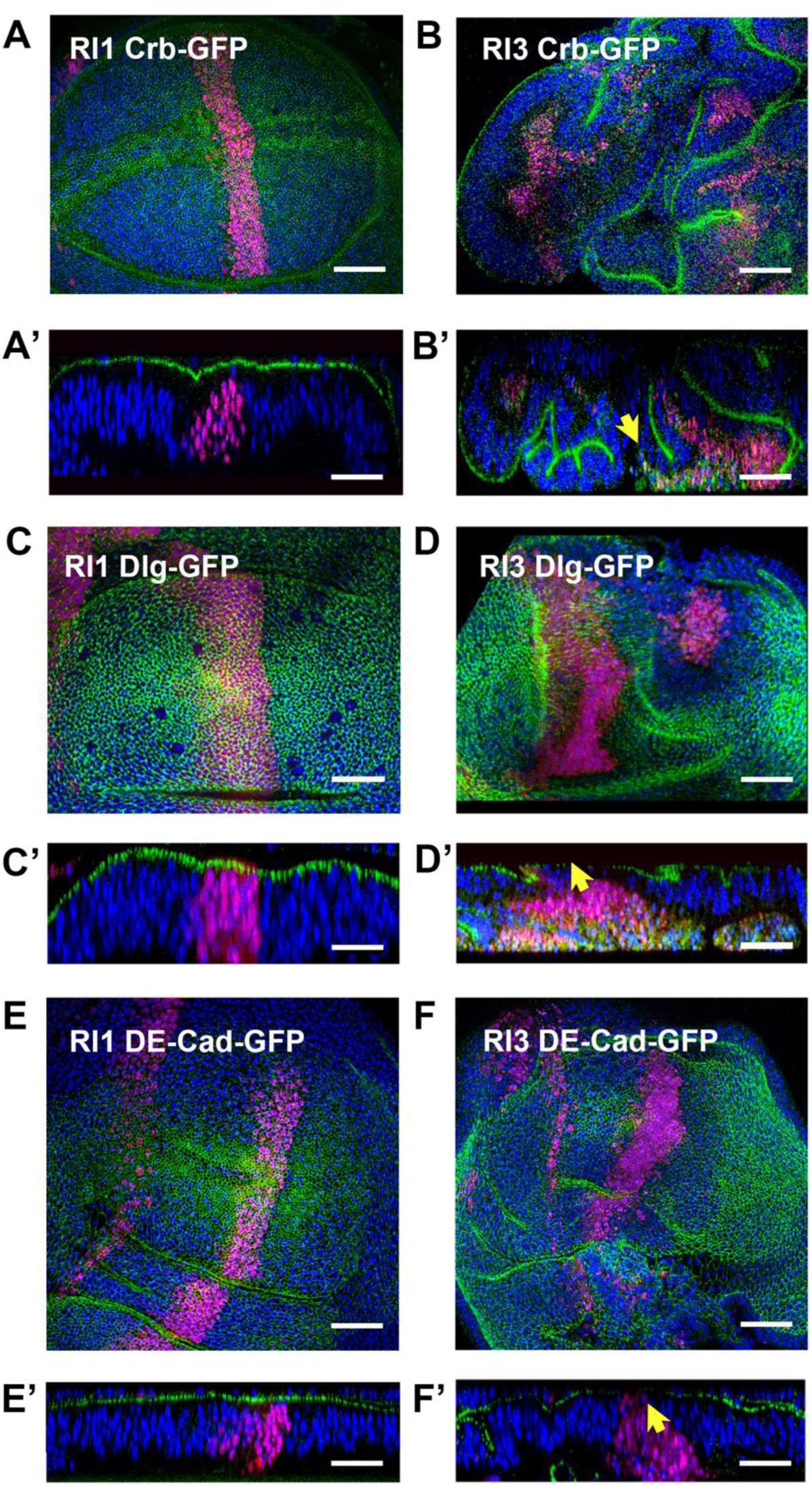
Repetitive regeneration induction leads to the loss of apico basal polarity. **A** and **B)** Crumbs (Crb) expression (green) in RI1 and RI3 wing imaginal discs (frontal views). **A’** and **B’** display perpendicular cross-sections. Ptc expression in magenta and DAPI in blue. Yellow arrow points to the loss of Crb apical expression in RI3 discs. Scale bar is 30 µm. **C** and **D)** Disc large (Dlg) expression (green) in RI1 and RI3 wing imaginal discs (frontal views). **C’** and **D’** display perpendicular cross-sections. Ptc expression in magenta and DAPI in blue. Yellow arrow points to the loss of Dlg apical expression in RI3 discs. Scale bar is 30 µm. **E** and **F)** DE-Cadherin (DE-Cad) expression (green) in RI1 and RI3 wing imaginal discs (frontal views). **E’** and **F’** display perpendicular cross-sections. Ptc expression in magenta and DAPI in blue. Yellow arrow points to the loss of DE-Cad apical expression in RI3 discs. Scale bar is 30 µm.

In summary, upon sequential death challenges, the regenerating blastema suffers a neoplastic transformation in which cells undergoing uncontrolled proliferation, lose their apicobasal polarity leading to a failure in epithelial integrity.

### Infiltration of neoplastic regenerating overgrowths

The degradation of the ECM has been associated to neoplastic overgrowths and it is a key factor for tumors’ ability to invade other tissues ^32^. Indeed, the experimental elimination of the basal lamina (ECM) (e.g., upon Matrix Metalloproteases (MMPs) overexpression) permits tumor cells to leave their tissues of origin and invade different organs ^33^. We observed, upon induction of sequential cell death, that the ECM unsheathing the wing disc (Collagen IV, visualized using Viking-GFP) ^34^ suffers multiple fissures, exposing the epithelial tissues to the hemolymph (compare **Figure 3A** to **3B**). As a consequence, circulating hemocytes, and those residing at the surface of the wing imaginal disc, become recruited and invade the imaginal epithelia, intermixing with the overgrowing neoplastic tissue (compare **Figure 3A’** and **3B’**). The degradation of the basal lamina associates to the ectopic expression of MMP1 (compare **Figure 3C** and **3D**).

**Figure 3.**
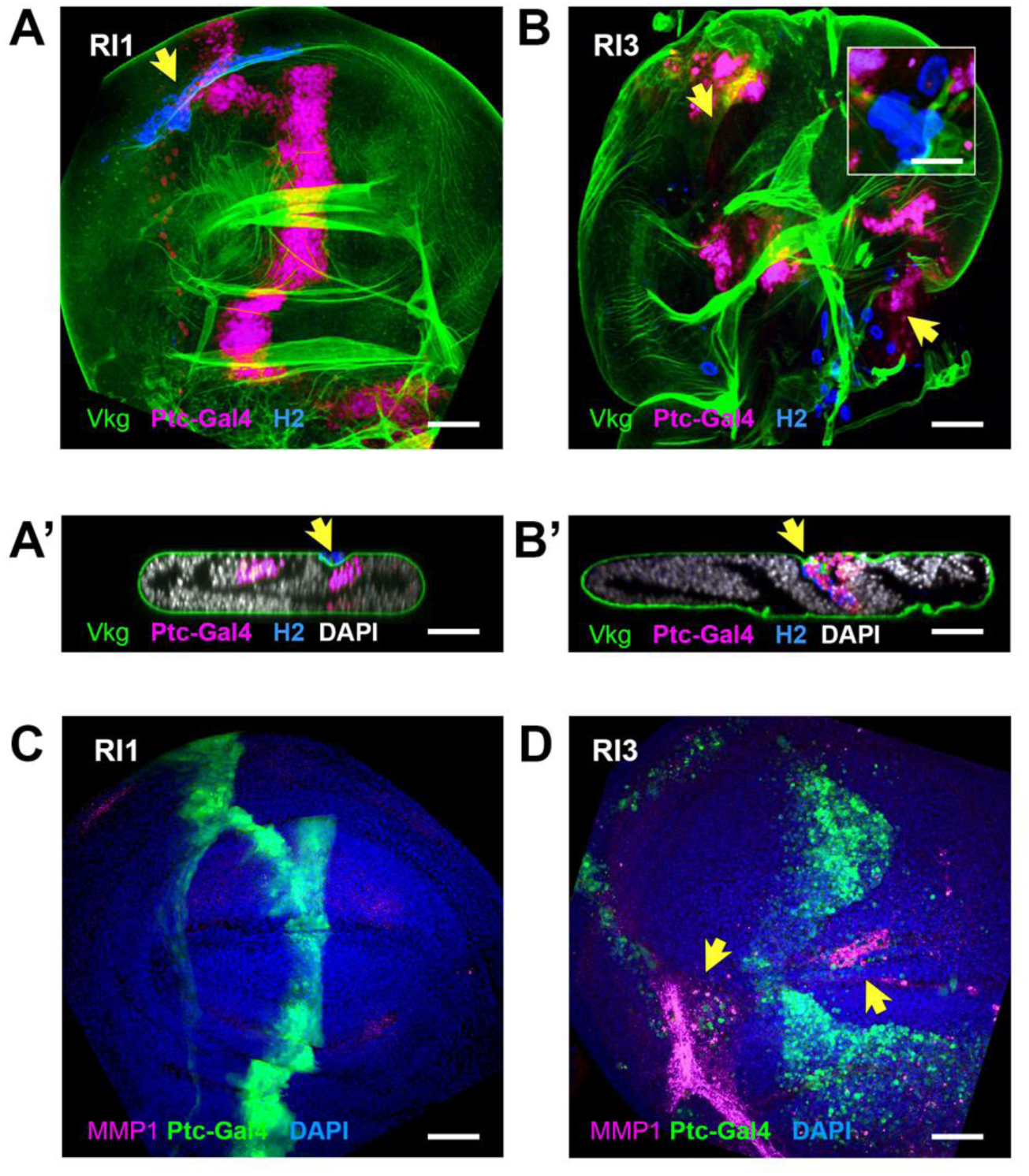
Repetitive regeneration induction leads to the break of the basal lamina and hemocytes invasion of the tumorigenic tissue. **A** and **B)** Viking-GFP (basal lamina) (green), Ptc/UAS-GFP (magenta) and H2 (hemocytes) in RI1 and RI3 wing imaginal discs (frontal views). Inset in **B** presents a high magnification detail of a hemocyte crossing a basal lamina gap. Scale bar is 10 µm. **A’** and **B’** display perpendicular cross-sections. Yellow arrow points to the loss of the basal lamina and hemocytes in RI3 discs. Scale bar is 40 µm. **C** and **D)** Matrix Metalloprotease 1-GFP (MMP1) expression (magenta) in RI1 and RI3 wing imaginal discs (frontal views). Ptc expression in green and DAPI in blue. Yellow arrow points to the ectopic expression of MMP1 in RI3 discs. Scale bar is 30 µm.

In *Drosophila*, embryonic hematopoiesis produces two types of hemocytes: phagocytic plasmatocytes and crystal cells ^35^. Both types persist into larval stages. The larva also possesses a hematopoietic organ, the lymph gland ^36^. Injury to the larva results in the release into circulation of a new type of hemocytes, the lamellocytes, from the lymph gland ^37^, which have been shown to participate in the formation of melanotic tumors. We do not know if hemocytes degrade the basal membrane or they are recruited to repair it. Yet, in humans, the presence of infiltrating immune cells in tumors correlates with cancer progression and bad prognosis ^38^, suggesting that their presence within the neoplastic regenerating blastema in our model may be a sign of malignancy.

### Migratory capability of neoplastic imaginal cells

Uncontrolled wing disc overgrowths generated upon sequential cell death insults also affect patterning. Several markers such as Cubitus Interruptus (Ci) and Hedgehog (Hh), which in the wild type show a precise complementary expression delimiting anterior and posterior compartments ^39^, partially intermingle at the anterior-posterior border (compare **Figure 1A** to **S1A’**). Further, the expression of Wingless (Wg) was also altered and their native expression at wing pouch edges and at the wing dorsoventral boundary ^40^ became enlarged and aberrant (compare **Figure S1B** to **S1B’**).

**Figure S1.**
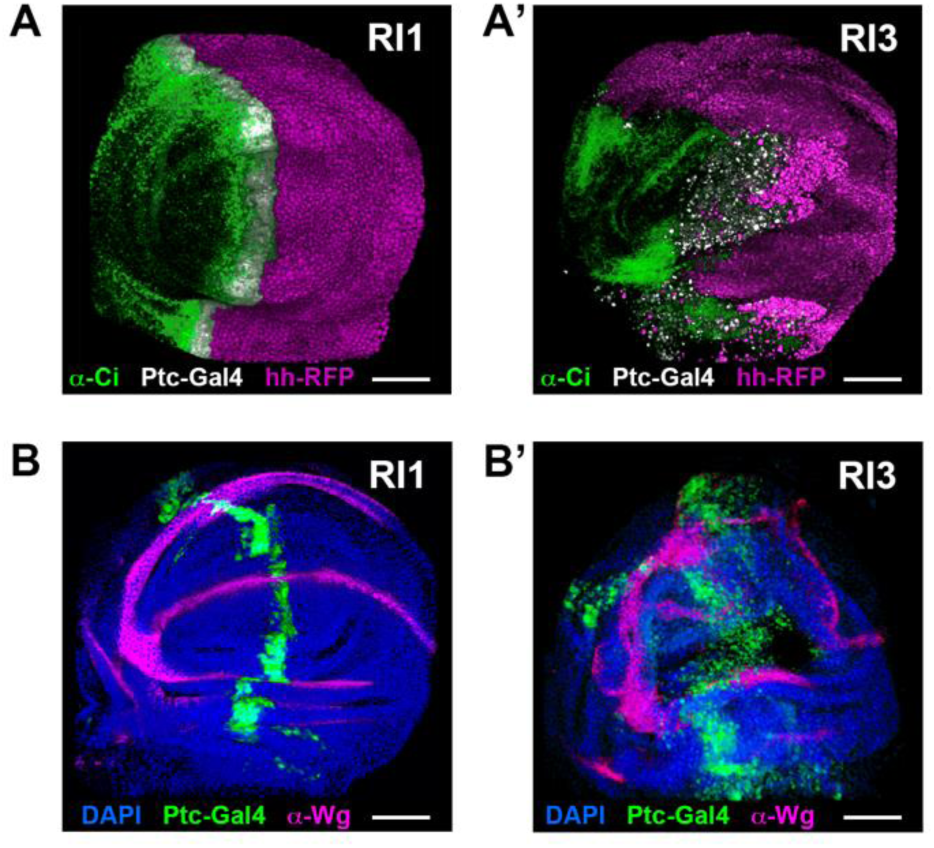
Patterning defects in wing imaginal discs after repetitive induction of Rpr. **A** and **A’)** Anterior (Ci) (green) and posterior (Hh-RFP) (magenta) compartment expression in RI1 (**A**) and RI3 (**A’**) wing discs. Ptc expression is shown in white. Scale bar is 30 µm. **B** and **B’)** Wingless (Wg) wing pouch expression (magenta) in RI1 (**B**) and RI3 (**B’**) wing discs. Ptc expression is shown in green and DAPI in blue. Scale bar is 30 µm.

The fading of compartment borders and the loss of epithelial integrity after successive rounds of cell death, advocate for a transition of regenerative neoplastic imaginal cells towards a mesenchymal fate with the acquisition of migratory capabilities. To analyze the migratory properties of death-targeted cells in the *ptc* domain, real time and lineage-traced expression was studied by G-TRACE (see **Experimental Procedures**). We observed, as expected, that after a single round of apoptosis/regeneration the *ptc* pattern was rebuild essentially as observed in wild type larvae (**Figure 4A**). Cells, once expressing *ptc*, but not anymore (RI1), were observed adjacent, mostly anterior, to the current pattern indicating that not all *ptc*-expressing cells were eliminated upon Rpr overexpression. In contrast, after 3 rounds of apoptosis/regeneration (RI3), we observed that the *ptc* pattern enlarged extensively (**Figures 4B**) and that a considerable number of cells that once expressed *ptc* did not die. Importantly, many of the cells surviving multiple rounds of death induction detach from the *ptc* domain turning into “walking dead cells” acquiring invasive properties and spreading to other territories (**Figure 4B’**). Elsewhere, non *ptc*-expressing cells get recruited to the neoplastic overgrown tissue, contributing to its heterogeneous character (**Figure 4B’’**).

**Figure 4.**
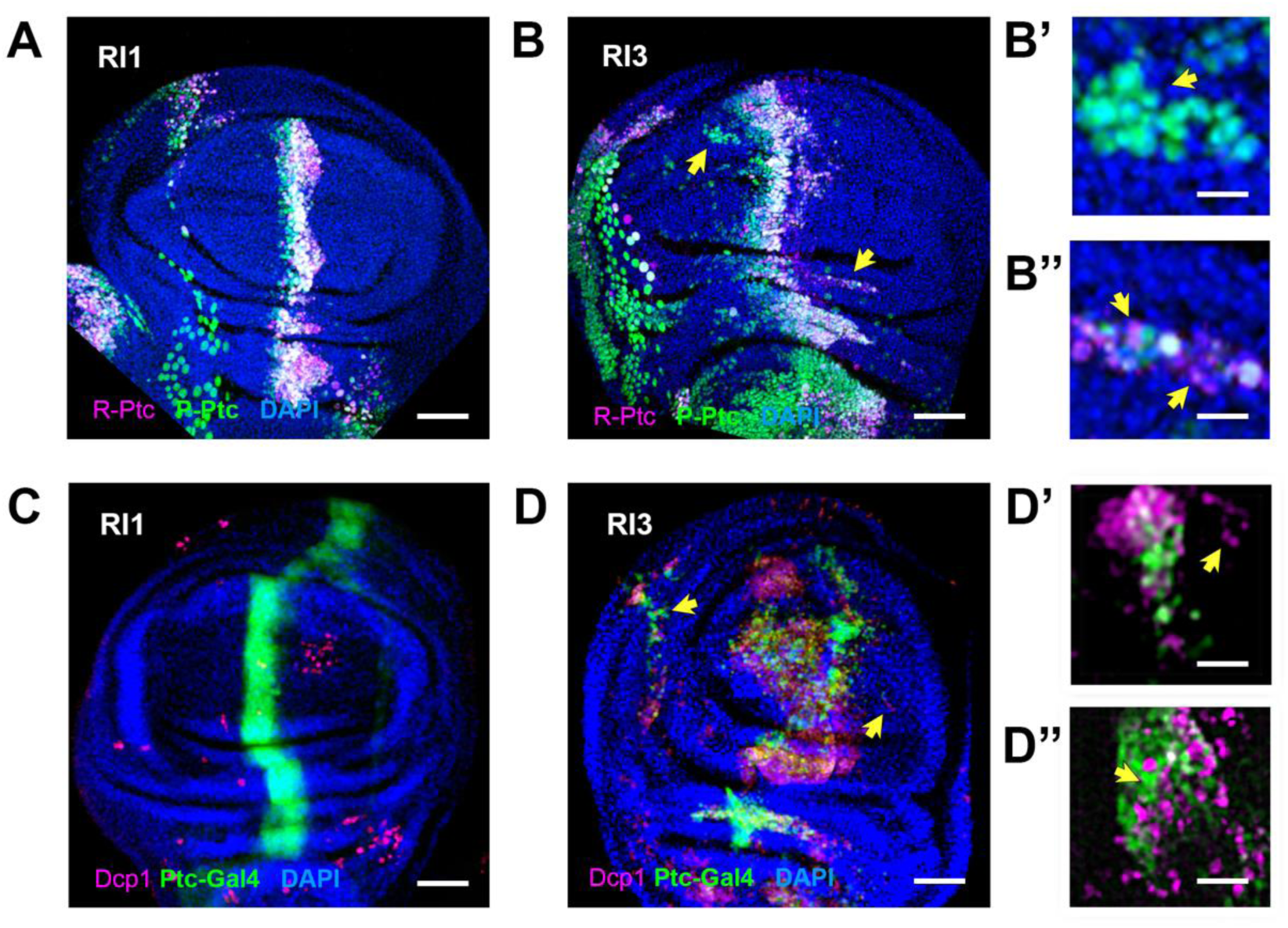
Repetitive regeneration induction leads to the break of the basal lamina and hemocytes invasion of the tumorigenic tissue. **A** and **B)** GTRACE analysis of Rpr-expressing cells after one (RI1) (**A**) and three rounds (RI3) (**B**) of apoptosis induction. All cells once expressing Ptc are permanently labelled in green (P- Ptc). Magenta depicts those cells expressing the Gal4 recently (R-Ptc) at the time of the analysis. DAPI expression is shown in blue. Arrows in **B** point to 1) “walking dead cells” detached from the Ptc domain (**B’**) and 2) non Ptc-expressing cells recruited to the overgrowing tissue (**B’’**). Scale bar is 30 µm in **A** and **B** and 5 µm in **B’** and **B’’**. **C** and **D)** Dcp1 (dead cells) expression (magenta) in RI1 (**C**) and RI3 (**D**) wing imaginal discs (frontal views). Ptc expression is shown in green and DAPI in blue. Yellow arrows point to 1) dying cells in areas outside the Ptc domain (**D’**) and 2) Dpc1-negative death-surviving Ptc-expressing cells (**D’’**). Scale bar is 30 µm in **C** and **D** and 5 µm in **D’** and **D’’**.

### Cell death in regenerating neoplastic overgrowths

The fact that many cells in the regenerating blastema appear to survive cell death even after suffering three rounds of death induction was puzzling. We monitored cell death by analyzing the activation of the effector caspase Dcp1 and found that, (compare **Figure 4C** to **4D)**, the majority of the *ptc*-expressing region is comprised of dying cells. In addition, some dying cells were observed in areas outside the *ptc* domain, which was indicative of non-autonomous death events or the presence of “walking death cells” (**Figure 4D’**). Further inspection of the *ptc* region revealed Dcp1 negative death-surviving *ptc*-expressing cells (**Figure 4D’’**) of tumorigenic potential.

### DNA damage control and centrosome instability

Correct DNA replication is a crucial event on the S phase of the cell cycle for the maintenance of genome integrity and accurate chromosome segregation is crucial to ensure cell viability. Remarkably, most cancers correlate with genomic instability as a result of failures on the DNA damage safeguard mechanisms and errors in mitosis are a frequent source of chromosomal alterations as those observed in cancer cells ^26,41^. Tumoral cells, in many cases, also display centrosome and spindle abnormalities, supernumerary centrosomes or even loss of centrosomes.

Our data indicated that a fraction of regenerating cells become refractory to death, proliferate in an uncontrolled manner and acquire migratory characteristics. We then asked if these cells avoiding apoptosis could, eventually, had suffered replication stress and DNA damage, or centrosome instability.

In the context of the anomalous epithelial regeneration caused by repeated death/regeneration events, either unrepaired DNA damage, leading to mutations triggering tumor-enabling factors or altered centrosomal segregation during mitosis ^42,43^, could result in the generation of malignant neoplastic overgrowths. To evaluate these possibilities, we studied the presence of DNA double-strand breaks (DSBs) by immunostaining for gamma H2A. Comparing control and experimental discs, we found that the number of cells with DNA damage increased significatively in overgrown discs (compare **Figure 5A1** to **5A2**), indicating that repetitive stress imposed on the regenerative blastema affects DNA repairing capabilities (**Figure 5B**). As a result, mutations may accumulate, letting the generation of tumorigenic cells. Importantly, we observed in this condition that some cells expressing apoptotic and DNA damage markers, were negative for *ptc*; while others, positive for *ptc* and for DNA damage markers were non-apoptotic (**Figure 5C**). Secondly, we evaluated centrosomal instability employing antibodies anti-γ Tubulin and anti-α Tubulin, to label the centrosomes and the mitotic spindle and microtubule organizing center respectively. Control discs showed stable bipolar spindles and a correct number and organization of their centrosomes (**Figure 5D1**). In contrast, experimental discs, carrying the UAS-Rpr transgene after sequential regeneration challenges, show, at quite high frequency (**Figure 5E**) centrosome and spindle aberrations (elongated, multipolar, unipolar or asymmetric polar spindles) (**Figure 5D2-5D4**), characteristic of cancer cells. Last, we evaluated the development of aneuploids after repeated series of apoptosis/regeneration cycles by flow cytometry. Histograms were retrieved for dissociated tissues, comparing the DNA content of cells from control wing discs challenged just one time to those of selected discs with overgrowths after 3 rounds of apoptosis/regeneration. The percentage of GFP positive cells was low for both samples as the *ptc* pattern covers just a small portion of the wing disc, although, obviously, it increased in the overgrown tissue samples. While more cells in G2 and M phases (4C) than in G1 (2C) were observed for both conditions, overgrown discs almost quadruplicate the average amount of aneuploid cells in controls. This increase was detected both for *ptc*-expressing (3, 8 X) and non-expressing cells (3,9 X) (compare **Figure 5F** to **5G**).

**Figure 5.**
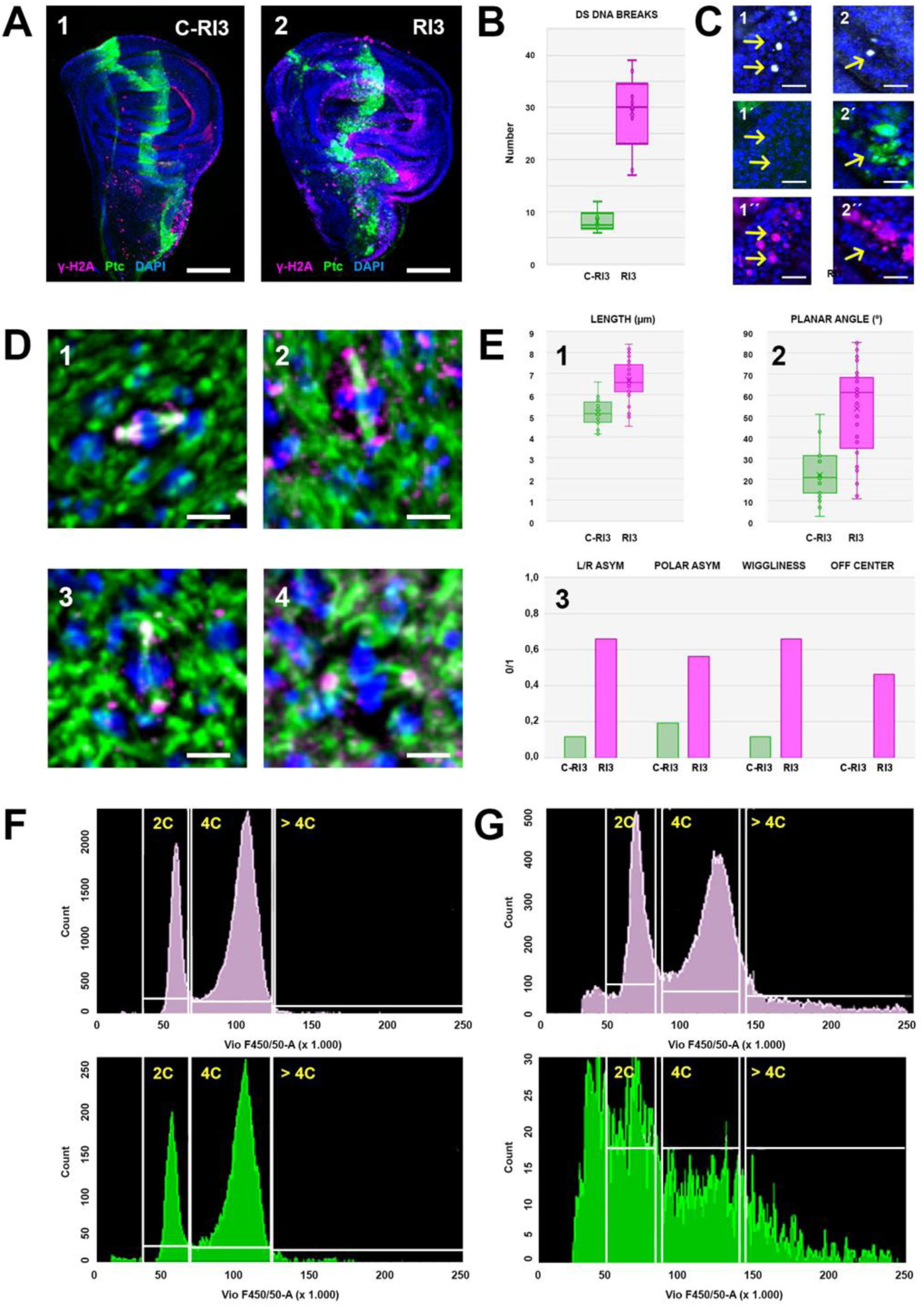
Sequential apoptosis induction affects DNA damage repair and chromosome stability. **A)** ã H2a labelling (magenta) of control RI3 (in the absence of Rpr) (**1**) and RI3 (**2**) wing imaginal disc depicting cells with double strand DNA breaks (DSBs). Green is Ptc / GFP and blue is DAPI. Scale bar is 50 µm. **B)** Quantification of the number of cells with DSBs on control RI3 (green) (n = 4) and RI3 (magenta) (n = 7) wing imaginal discs. The two-tailed P value between RI1 / Rpr and RI3 / Rpr equals 0.0017, which is very statistically significant. **C)** High magnification of overgrown areas of ã H2a (**1** and **2**) (white), Ptc / GFP (**1’** and **2’**) (green) and Dcp1 (**1’’** and **2’’**) (magenta) labelled RI3 wing discs. Arrows point to DNA damaged cells not expressing Ptc but dying (**1**, **1’** and **1’’**) and DNA damaged cells expressing Ptc but not dying (**2**, **2’** and **2’’**). **D)** Individual examples of spindle aberrations at metaphase in RI3 wing discs. (1) stable bipolar wild type spindle; (2) elongated; (3) multipolar; and (4) asymmetric spindles. Green is α Tubulin (spindles), magenta is γ Tubulin (centrosomes) and blue is DAPI. Scale bar is 5 µm. **E)** Quantitative comparative analysis of numbers and frequencies of observed spindle aberrations at metaphase on control RI3 (green) (n = 26) and RI3 (magenta) (n = 41) wing discs. (**1**) length in µm. The two-tailed P value between control RI3 and RI3 / Rpr equals < 0.0001, which is extremely statistically significant.; (**2**) planar orientation in (°) The two-tailed P value between control RI3 and RI3 / Rpr equals < 0.0043, which is very statistically significant.; and (**3**) 0/1 of observed spindle aberrations: Left / Right spindle asymmetry, polar (end to end) asymmetry, wiggliness, and off-centered centrosomes. **F)** DNA histograms obtained by flow cytometry (FACS) of control RI3 wing discs cells. The first peak in each histogram corresponds to those cells with 2C DNA content and the second peak to cells with 4C DNA content. Cells with a DNA content bigger than 4C are on the right. In magenta are shown all cells and in green those corresponding to the Ptc expressing cells. **G)** DNA histograms obtained by flow cytometry (FACS) of RI3 wing discs cells. Data are represented as in **F**.

In summary, repetitive cell death induction in a regenerative blastema affects to different degrees the homeostasis of targeted cells. Some resist to death, but are prompt to non-repaired DNA damage. Others fail to undergo correct divisions, which results in aberrant chromosomal segregation, errors in cytokinesis or in the progression through M phase, and aneuploidies. The epithelial tissue (the wing disc), as a whole, responds to these local damages by stochastically and non-autonomously disturbing proliferation, death and DNA repair in nearby cells.

### Gene expression differences between the regenerating and the tumorigenic blastema

The presence of cells with heterogeneous behaviors in response to repeated death challenges may be explained by the emergence of distinct new cell subpopulations. Landscapes of pattern formation, proliferation, and growth at the single-cell level in wild-type wing imaginal discs have been previously characterized. Different topographical domains of these discs can be precisely distinguished using single-cell RNA sequencing (scRNA-Seq) analyses, which define specific cell cohorts into spatial subregions ^44–46^. Secondary clustering within each subregion facilitates analysis of patterning refinement.

To investigate the composition of cell populations in tumorigenic and regenerative wing discs, we conducted a systematic analysis of single-nucleus transcriptomic datasets (snRNA-Seq). Data were generated from dissociated experimental discs and their respective controls using the 10X Genomics Single Cell platform. We analyzed pooled samples comprising 50 individually selected imaginal discs per condition: Regeneration Control (RC), Regeneration (R), Tumorigenesis Control (TC), and Tumorigenesis (T) (see **Experimental Procedures**). Although the total number of cells captured slightly varied between samples, the median number of genes detected per cell remained consistent at approximately 900 genes per cell, and the median unique molecular identifier (UMI) counts per cell were stable at around 2.000 per cell. To annotate individual cell clusters, we used the databases generated by Deng et al ^45^ as reference.

Applying UMAP (Uniform Manifold Approximation and Projection) dimension reduction ^47^ to data from Regeneration control discs, we distinguished distinct myoblasts (DIR and IND) and tracheal cells (TRACHEA) clusters, unambiguously marked by the expression of genes such as *twist* (*twi*) and *Mef2* ^48,49^ or *breathless* (*btl*) ^50,51^. Conversely, the majority of nuclei (n = 39.113) from the wing disc epithelia formed a continuum without clear-cut clustering boundaries. To distinguish epithelial subdomains, we employed the combinatorial expression of 40 binarized landmark genes ^45^. This spatial reference map allowed us to assign epithelial cells with high confidence to four regions: the wing pouch (POUCH), hinge (HINGE), notum (NOTUM), and peripodial epithelium (PE) (**Figure 6A**). This distribution ^45^ recapitulates proximodistal axis (PD) marker gene expression patterns. These general domains were further subdivided into a total of 30 subclusters (**Figure 6B**).

**Figure 6.**
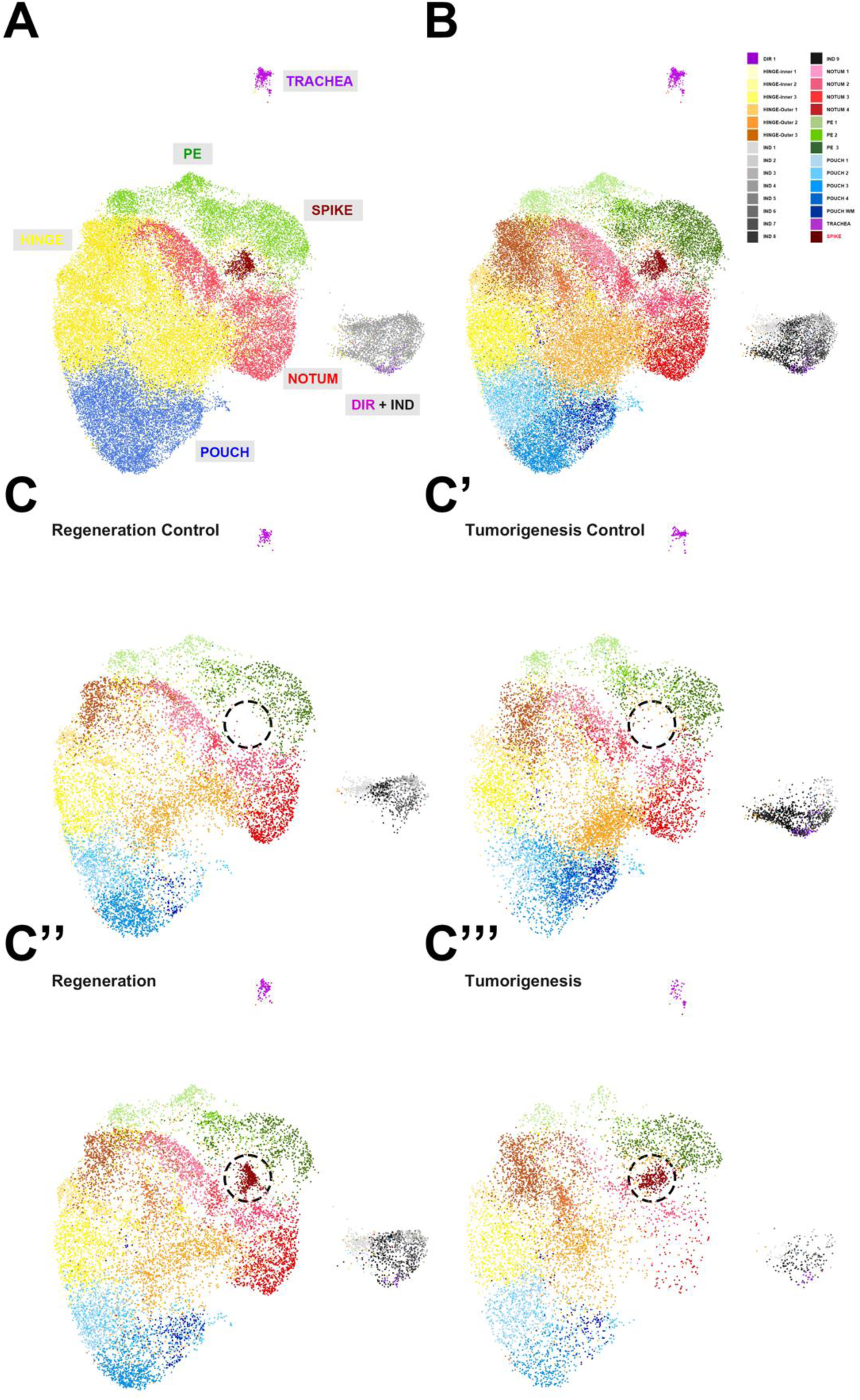
A single-cell survey of transcriptomic dynamics comparing regeneration and tumorigenesis conditions. **A)** Accumulated UMAP embedding (considering controls and experimental conditions altogether) of annotated major cell types corresponding to the notum, hinge, pouch, peripodial epithelium (PE), trachea, direct (DIR) and indirect (IND) muscles plus an undescribed population (SPIKE). **B)** UMAP embedding showing 30 further defined subclusters. **C-C’’’)** Cell-type composition across conditions: regeneration control **(C)**, tumorigenesis control **(C’)**, regeneration **(C’’)** and tumorigenesis **(C’’’)**. The SPIKE domain is encircled by a discontinued circumference.

When comparing data from the different experimental conditions, Regeneration and Tumorigenesis to their respective controls, we unexpectedly identified a cell population, that we termed SPIKE, that did not align with any of the described categories and was absent or strongly reduced in controls. This population exhibited a unique UMI composition and was predominantly distributed in between the NOTUM and PE domains (**Figure 6C** and **Figure S2**). SPIKE resembled cell populations (Blastemas 1 and 2) recently identified in regenerating wing imaginal discs subjected to the overexpression of the pro-apoptotic TNF orthologue Eiger in the wing pouch ^52^. Blastema 1 and 2 arise at the regenerating pouch/hinge interface and probably represent cells at two distinct steps of the regeneration process. Both share high expression levels for several genes including Ilp8, MMP1, Wg and Ets21C and are probably functionally equivalent to SPIKE (**Figure S3**).

SPIKE, despite communalities, displays a differential pattern of gene expression in regenerative and tumorigenic discs. A tumorigenic signature arises (**Figure 7**) displaying differential expression levels for several gene families, signaling elements or structural proteins (e.g. heat shock proteins are preferentially expressed in regenerative SPIKE cells, while members of the JNK signaling cascade or downstream targets, which appear to be a signature of SPIKE, are enriched in tumorigenic ones (see **Tables 1** and **2)**. JNK targets or genes indirectly targeted by JNK signaling, such as *MMP1*, *kay*, *puc*, *Socs36E*, *mys*, *zip* or *Rac2* are augmented both in the regenerating and tumorigenic blastemas. Importantly, upstream regulators of the JNK cascade such as *wnd* and *RhoGap18B* are enriched only upon tumorigenic induction, possible being instrumental on the increment of JNK activity detected in the blastema in this condition (see below **Figures 8A** and **8B**). Thus, when subjected to sequential cycles of cell death stimulation, failure of DNA and protein proofreading and JNK signaling misregulation, stand as potential contributors to cells transdetermination from a regeneration-committed state to an uncontrolled proliferative condition and, eventually, as causal factors of tumorigenic transformation.

**Figure 7.**
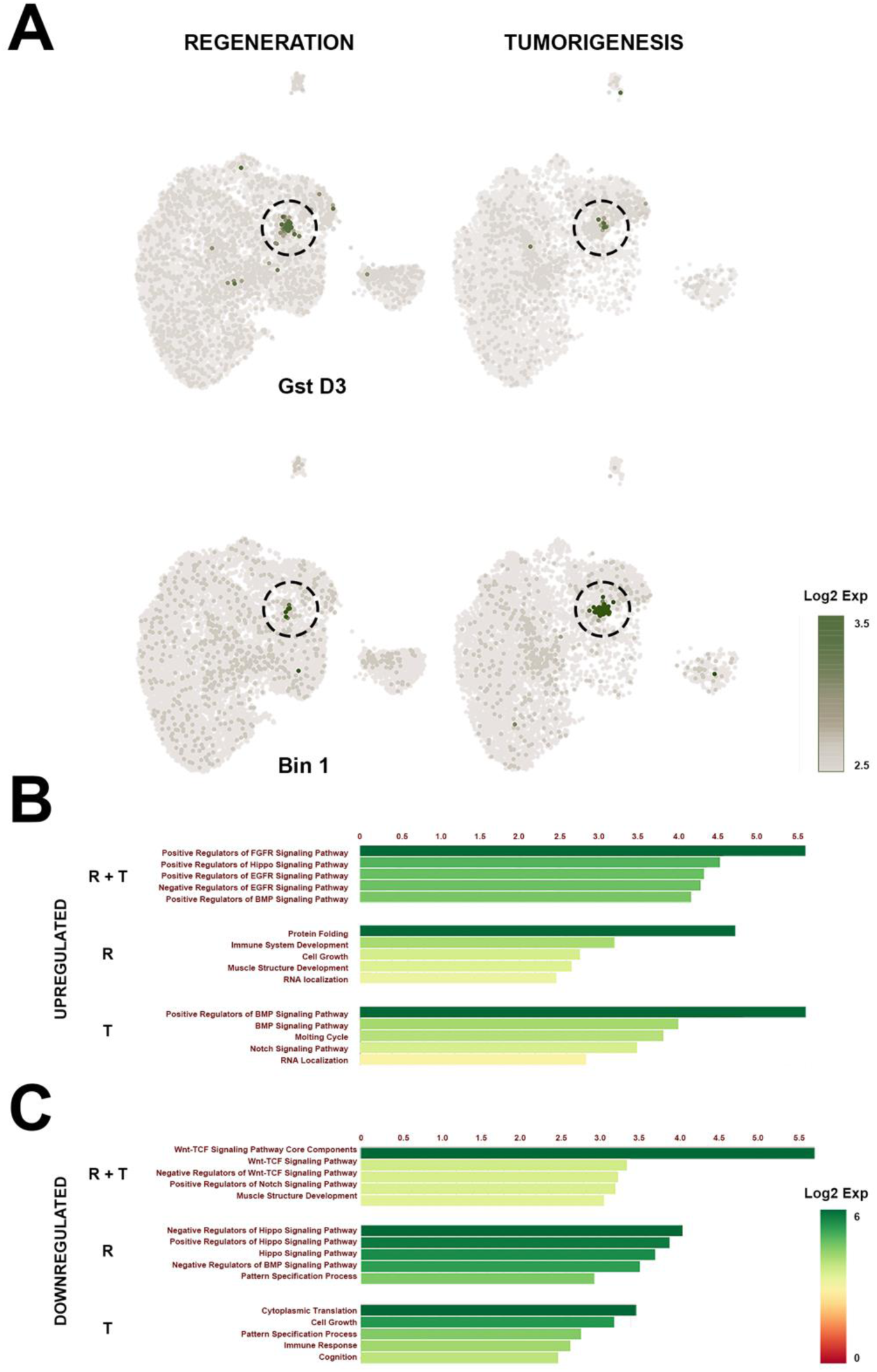
Differential gene enrichment of the regenerating and tumorigenic SPIKE (blastema) subcluster. **A)** Representative expression of differentially expressed genes in the SPIKE subcluster. *GstD3* is mostly enriched in the regenerating SPIKE, while *Bin1* is overrepresented in the tumorigenic SPIKE. **B)** Overrepresented (Log2 fold Expression) signaling and biological activity categories in the regenerative and tumorigenic (R + T) SPIKE or exclusively during regeneration (R) or tumorigenesis (T). **C)** Underrepresented categories as in **B**.

**Figure 8.**
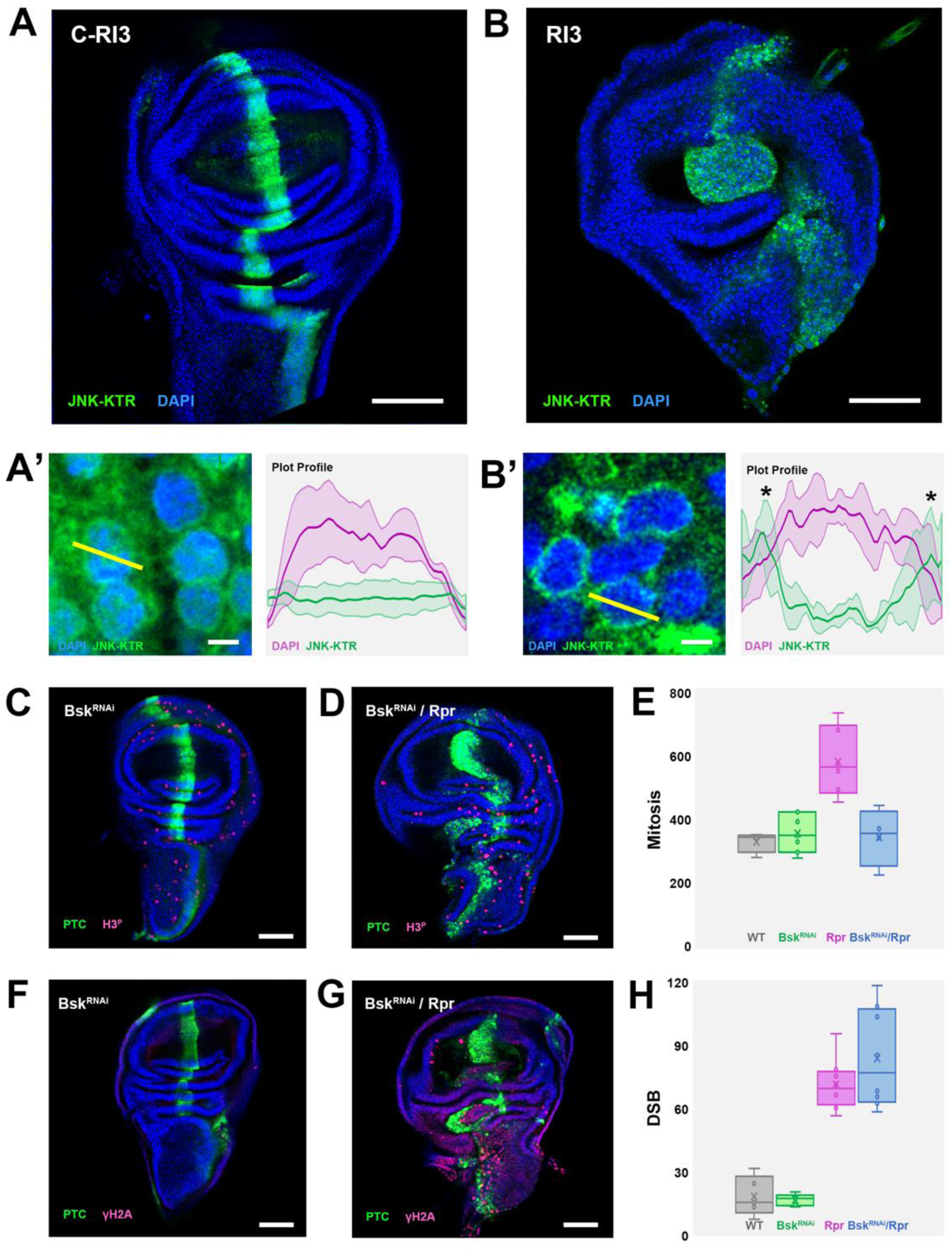
JNK signaling hyperactivation after repetitive regeneration induction leads to blastema overproliferation. **A** and **B)** JNK activity analysis of Rpr expressing cells employing a JNK-KTR sensor (green) on control RI3 (**A**) and RI3 wing discs (**B**). DAPI is blue. (**A’** and **B’**) High magnification images illustrating the subcellular distribution of the sensor. Yellow bars across cells are represented as plot profiles on the right of each panel. DAPI profile is shown in green and sensor profile is shown in magenta. Stars point to the increased cytoplasmic sensor localization observed in RI3 discs (**B’**). **C** and **D)** H3^P^ labelling (magenta) of a RI3 wing disc depicting cells in mitosis upon expressing a Bsk^RNAi^ transgene (**C**) or after dual overexpression of Bsk^RNAi^ and Rpr (**D**). Green is Ptc / GFP and blue is DAPI. Scale bar is 50 µm. **E)** Quantification of mitotic cells in wild type (grey) (n =4), RI3 / Bsk^RNAi^ (green) (n = 7), RI3 / Rpr (magenta) (n = 7) and RI3 / Bsk^RNAi^ + Rpr (blue) (n = 5) wing discs. The two-tailed P value between RI3 / Rpr and RI3 / Bsk^RNAi^ + Rpr equals 0.0035, which is very statistically significant. **F** and **G)** γ H2a labelling (magenta) of a RI3 wing disc depicting cells with DSBs after expressing a Bsk^RNAi^ transgene (**F**) or after dual overexpression of Bsk^RNAi^ and Rpr (**G**). Green is Ptc / GFP and blue is DAPI. Scale bar is 50 µm. **H)** Quantification of cells presenting DSBs in wild type (grey) (n = 4), RI3 / Bsk^RNAi^ (green) (n = 5), RI3 / Rpr (magenta) (n = 7) and RI3 / Bsk^RNAi^ + Rpr (blue) (n = 8) wing discs. The two- tailed P value between RI3 / Rpr and RI3 / Bsk^RNAi^ + Rpr equals 0.0719, which is not statistically significant.

**Table 1.**
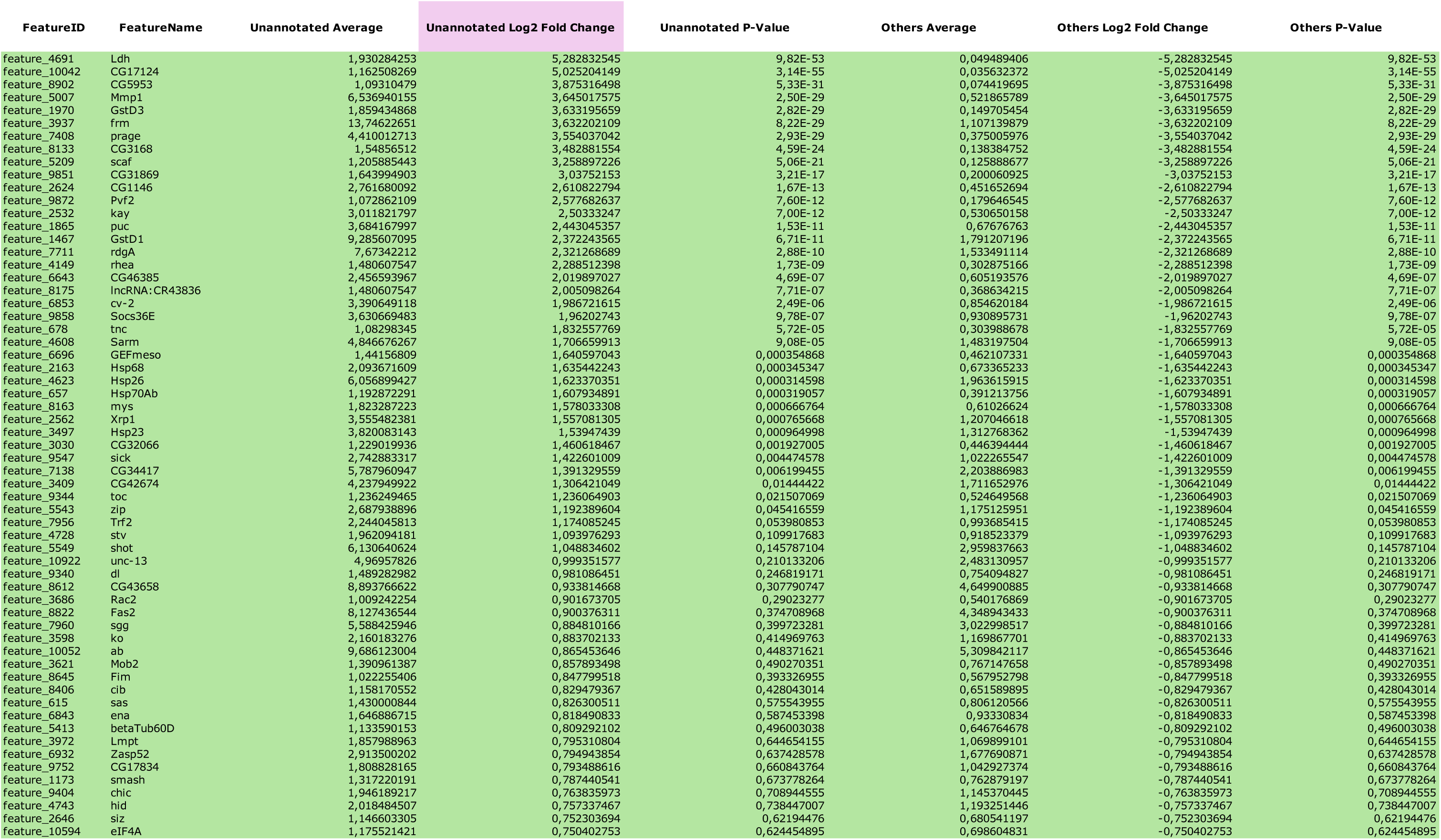
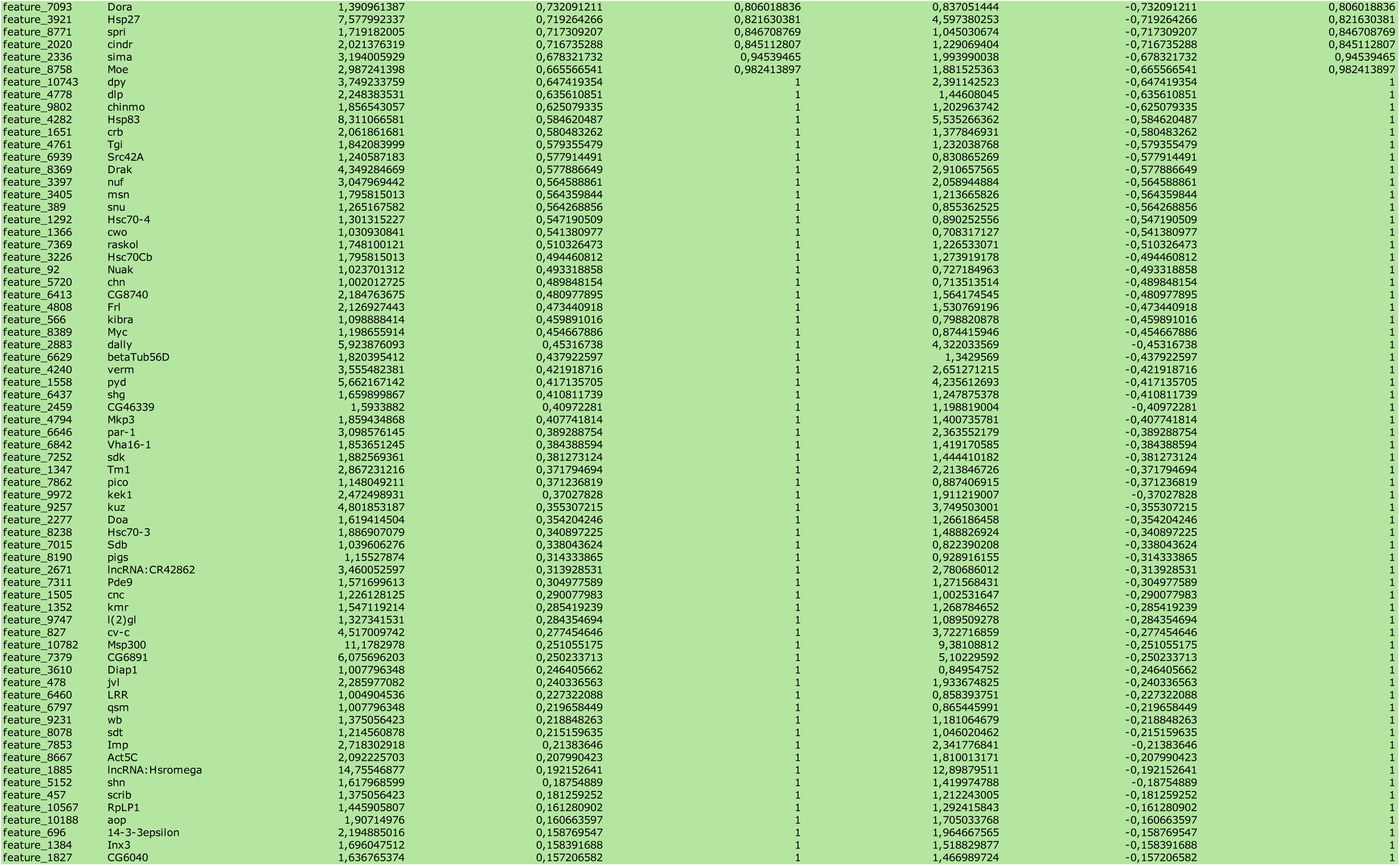
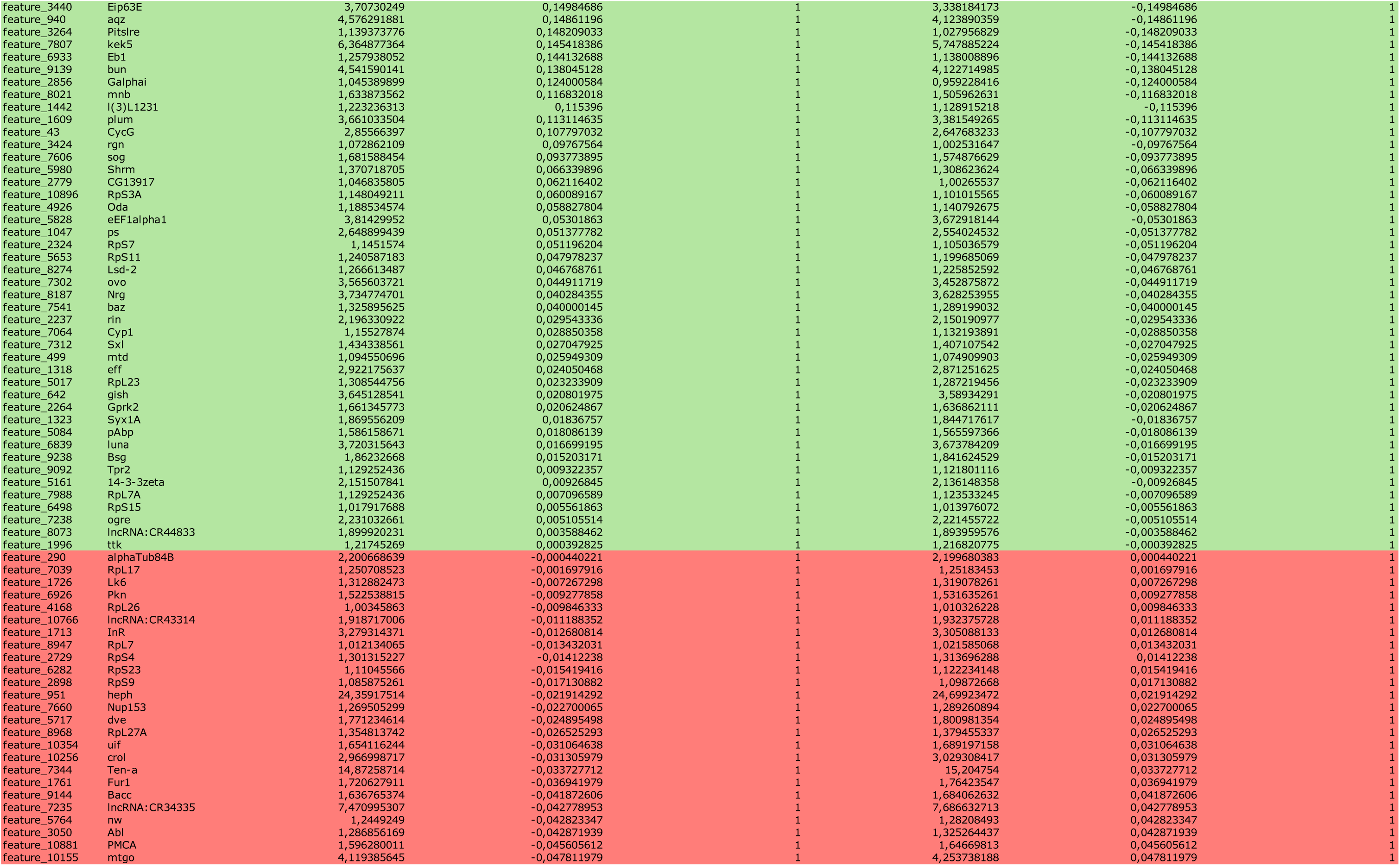
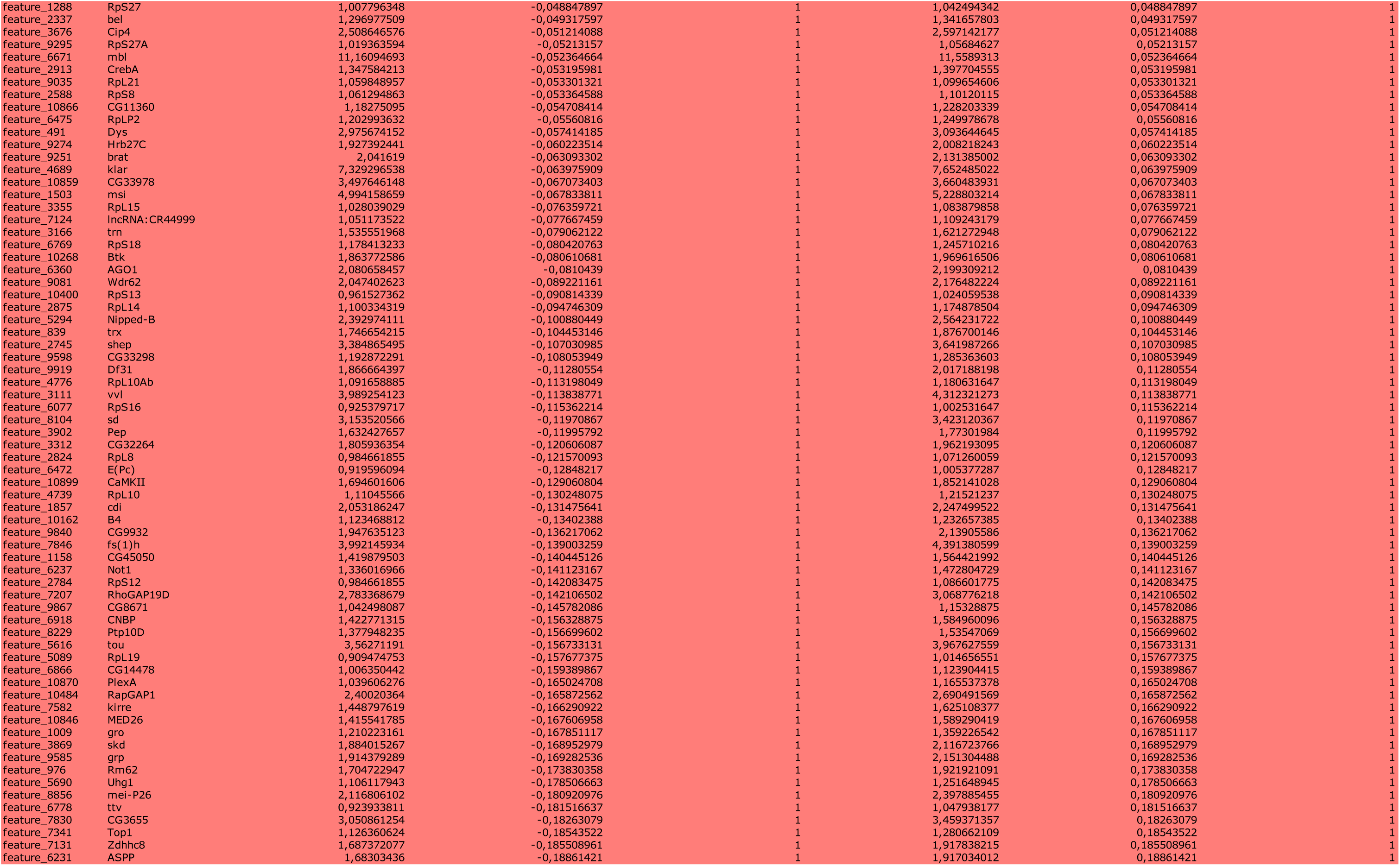
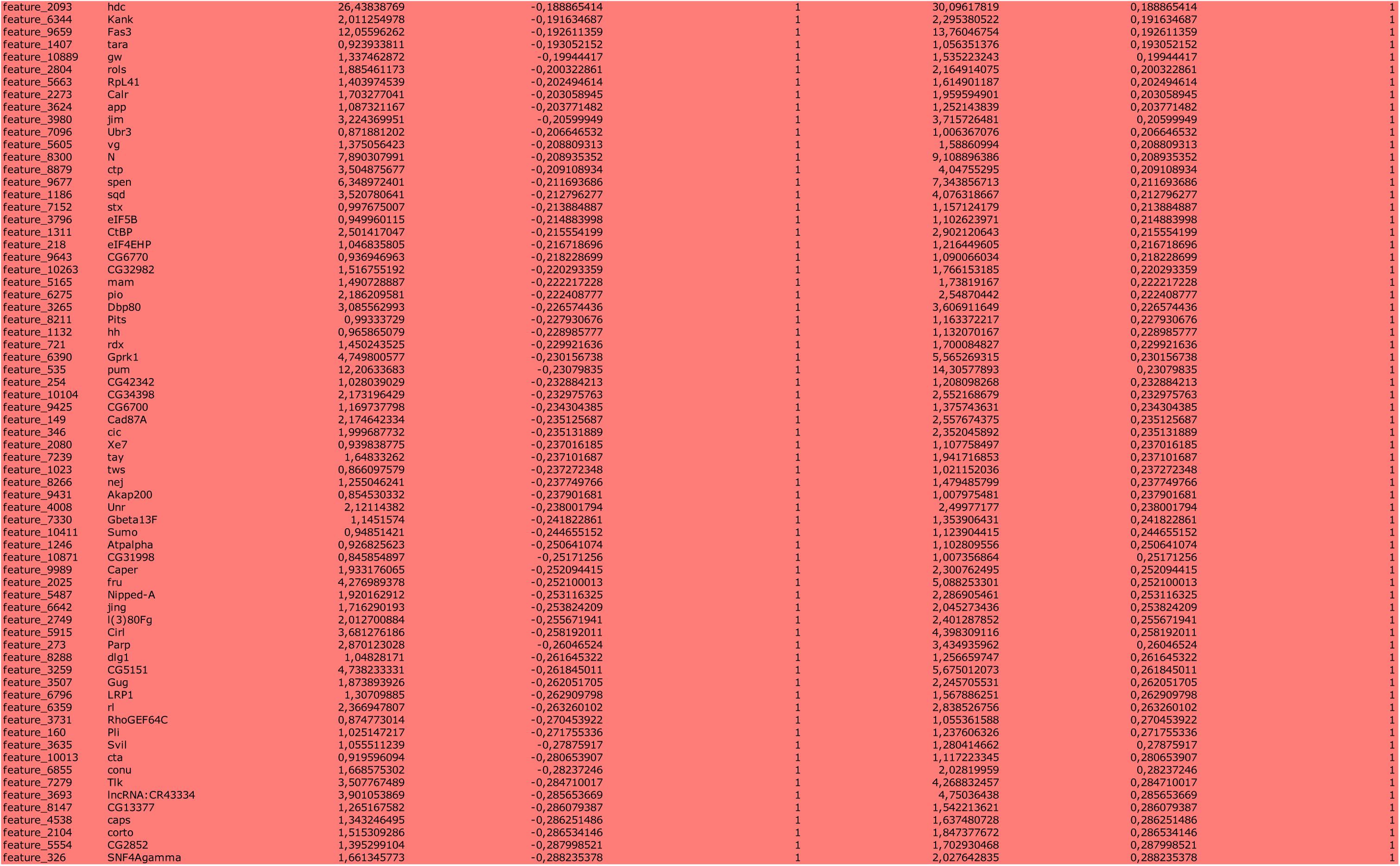
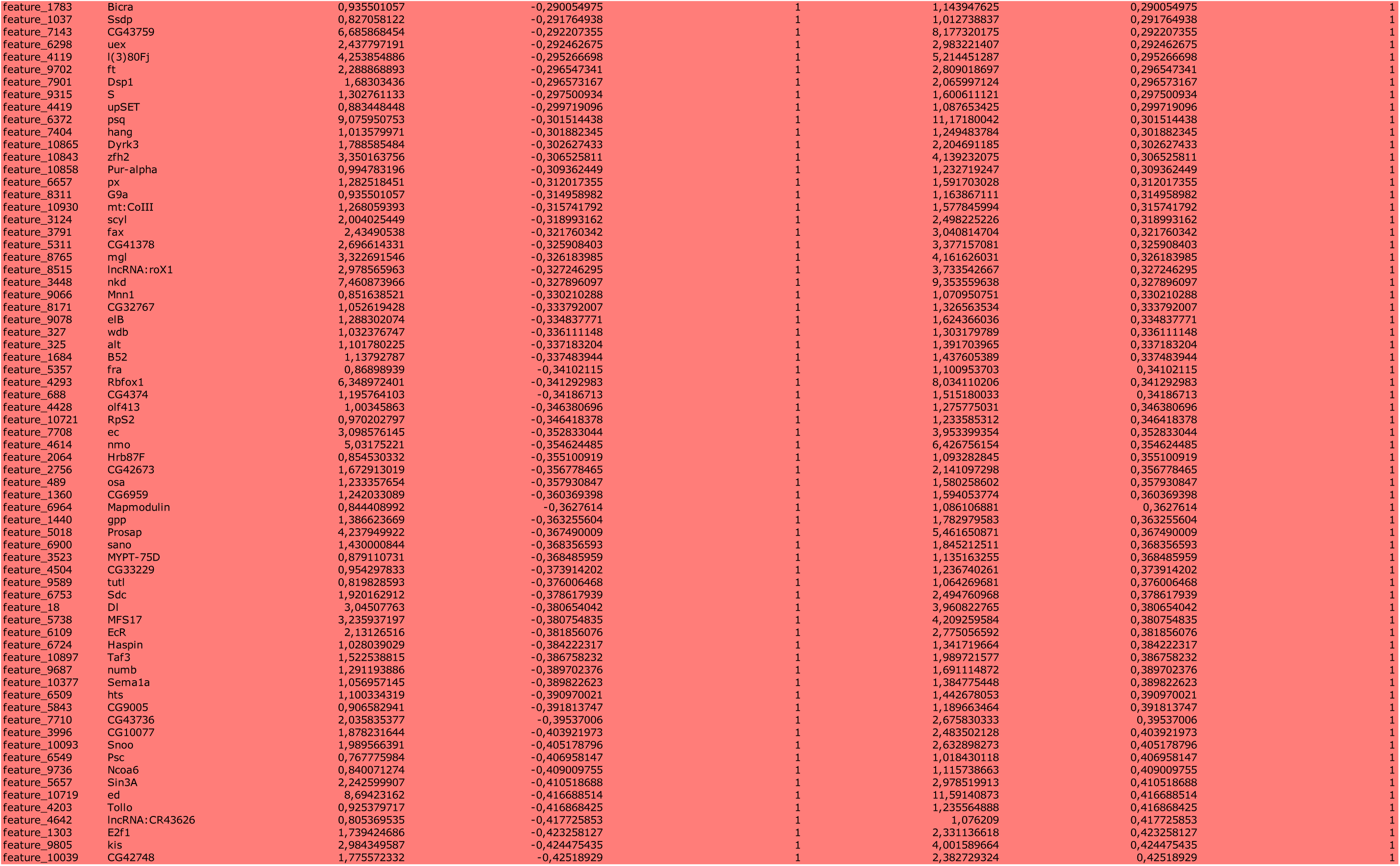
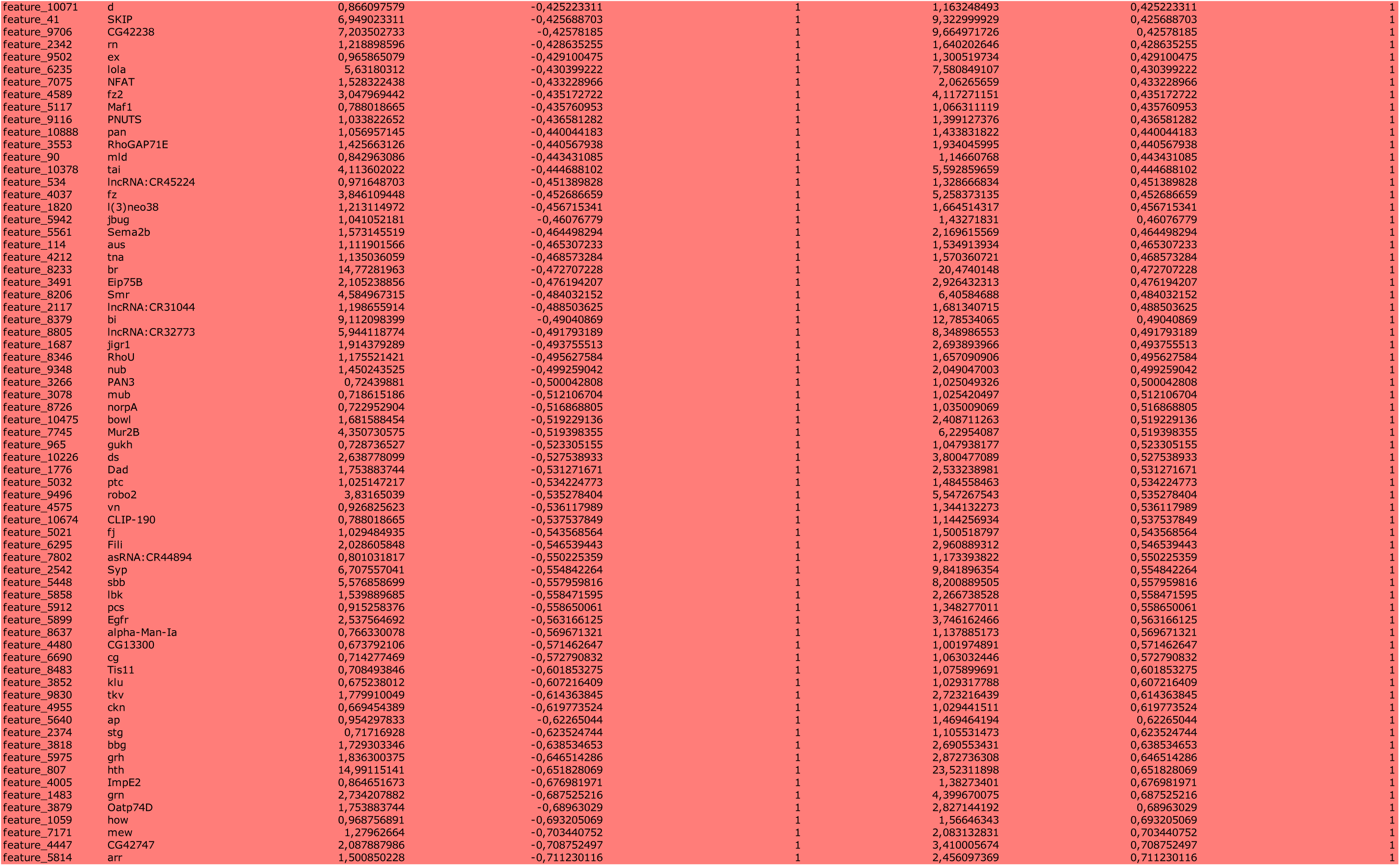
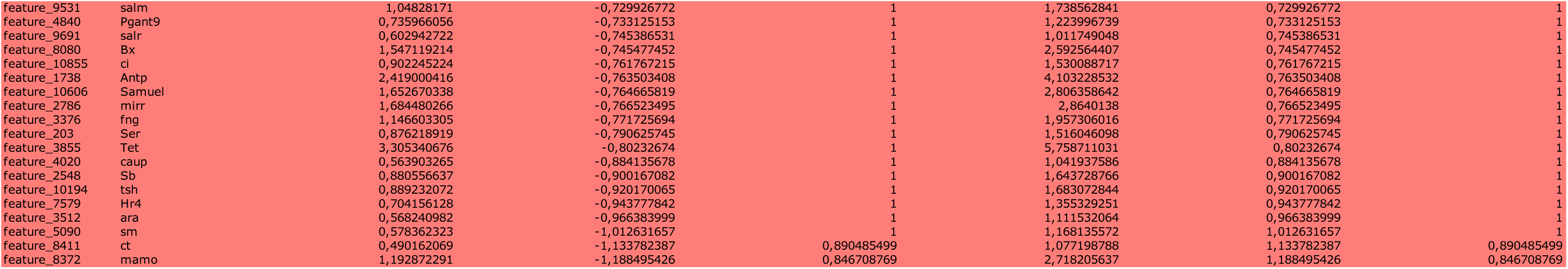
Differentially enriched or diminished expressed genes in the SPIKE subpopulation for Regenerating conditions.

**Table 2.**
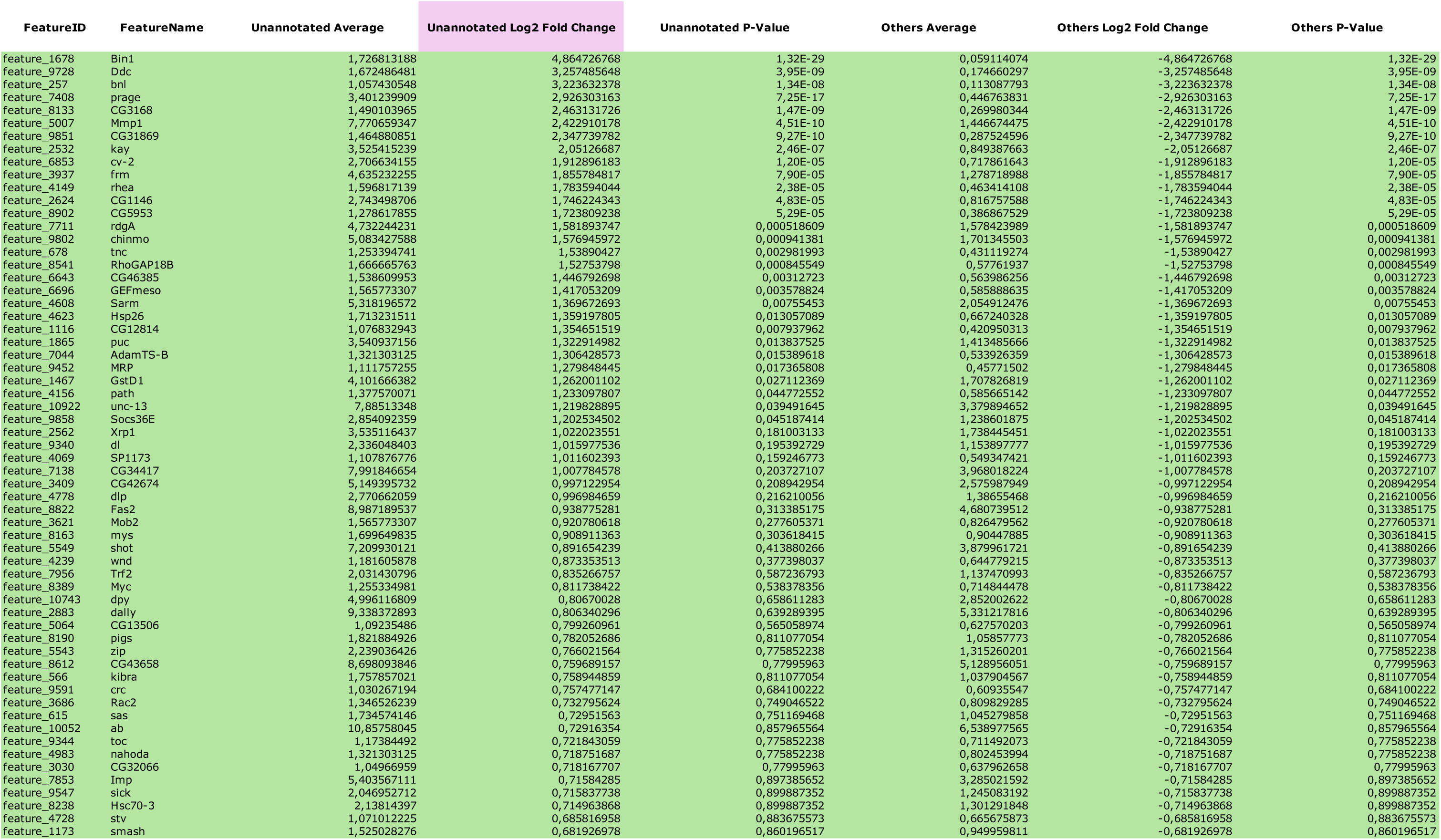
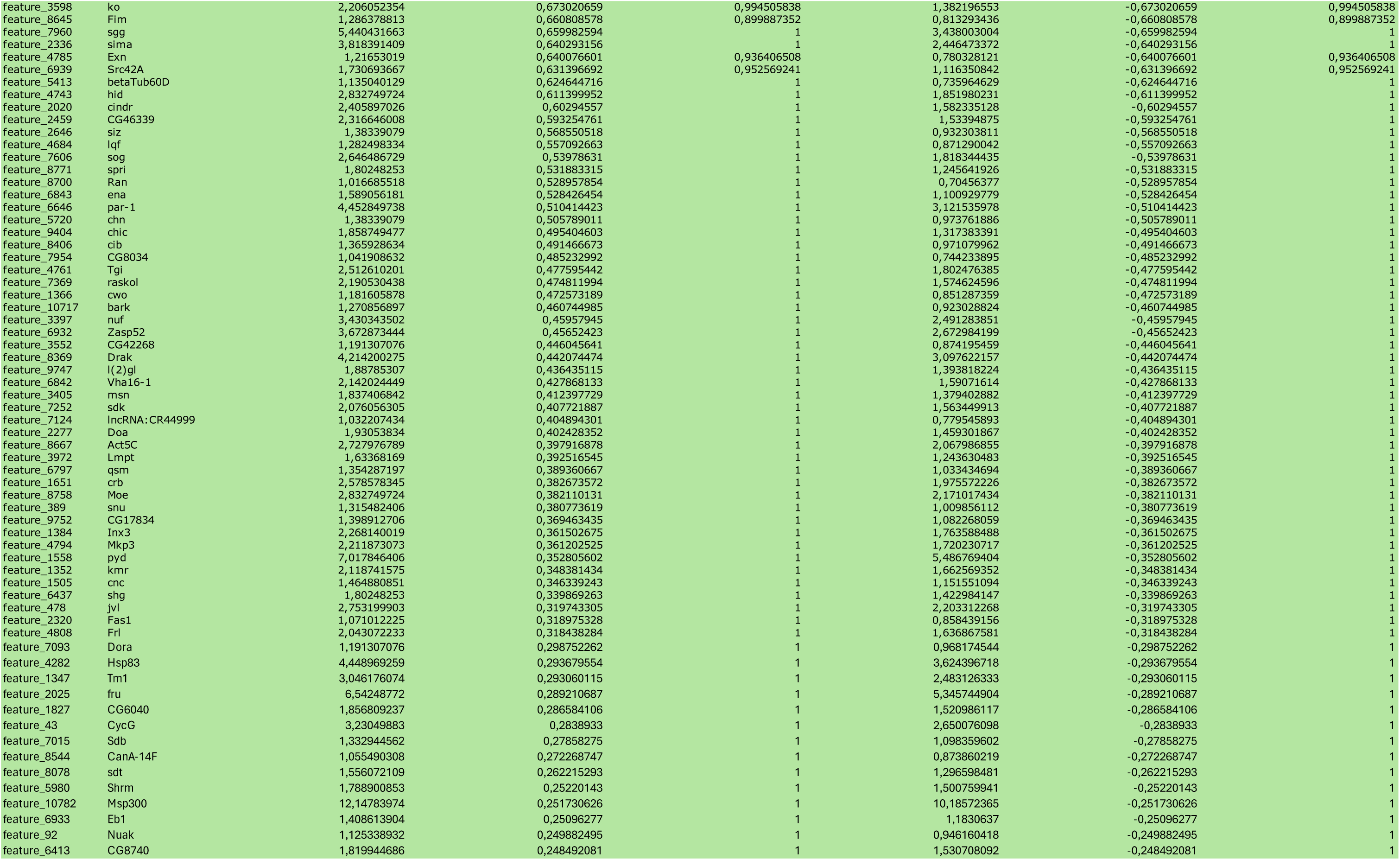
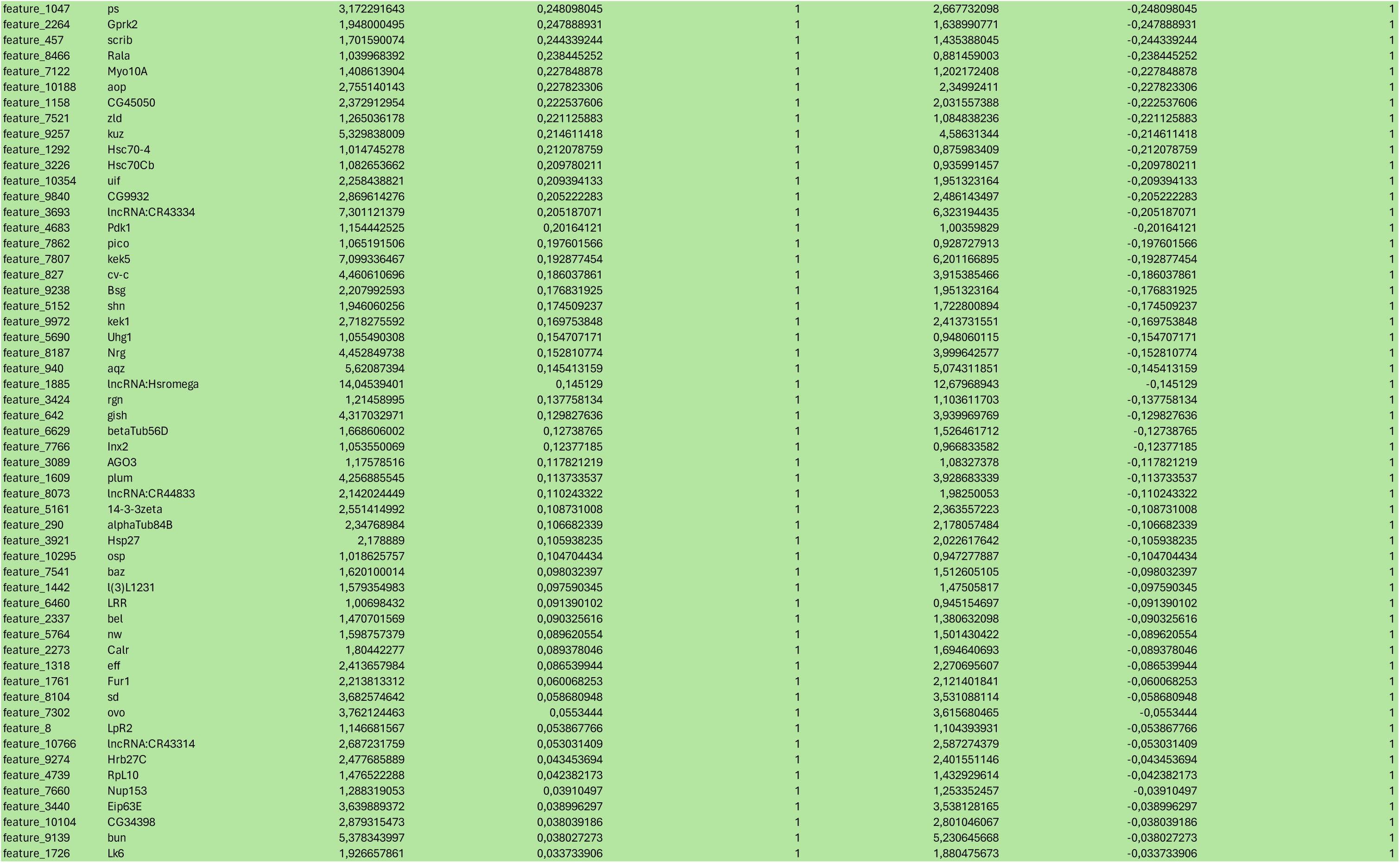
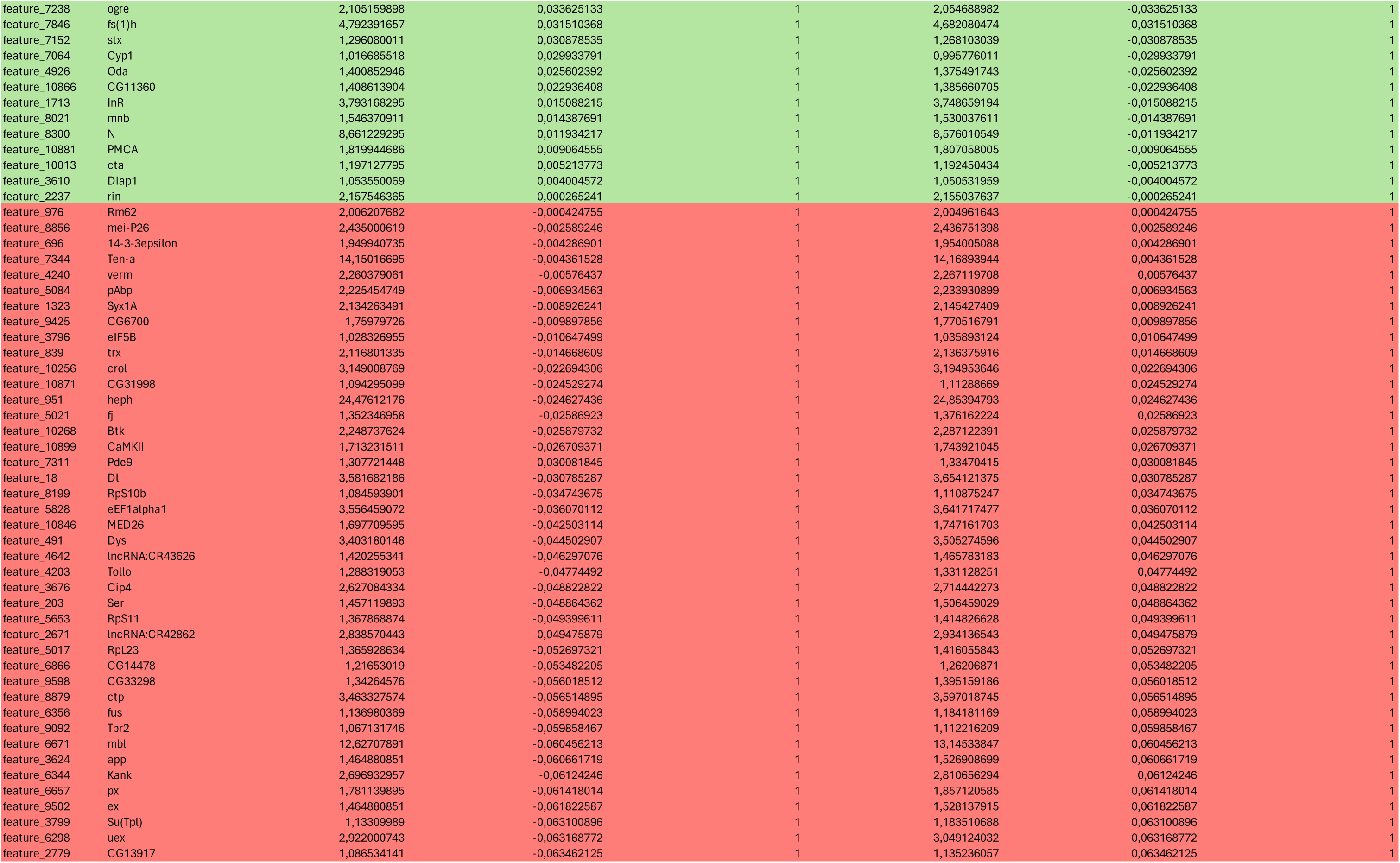
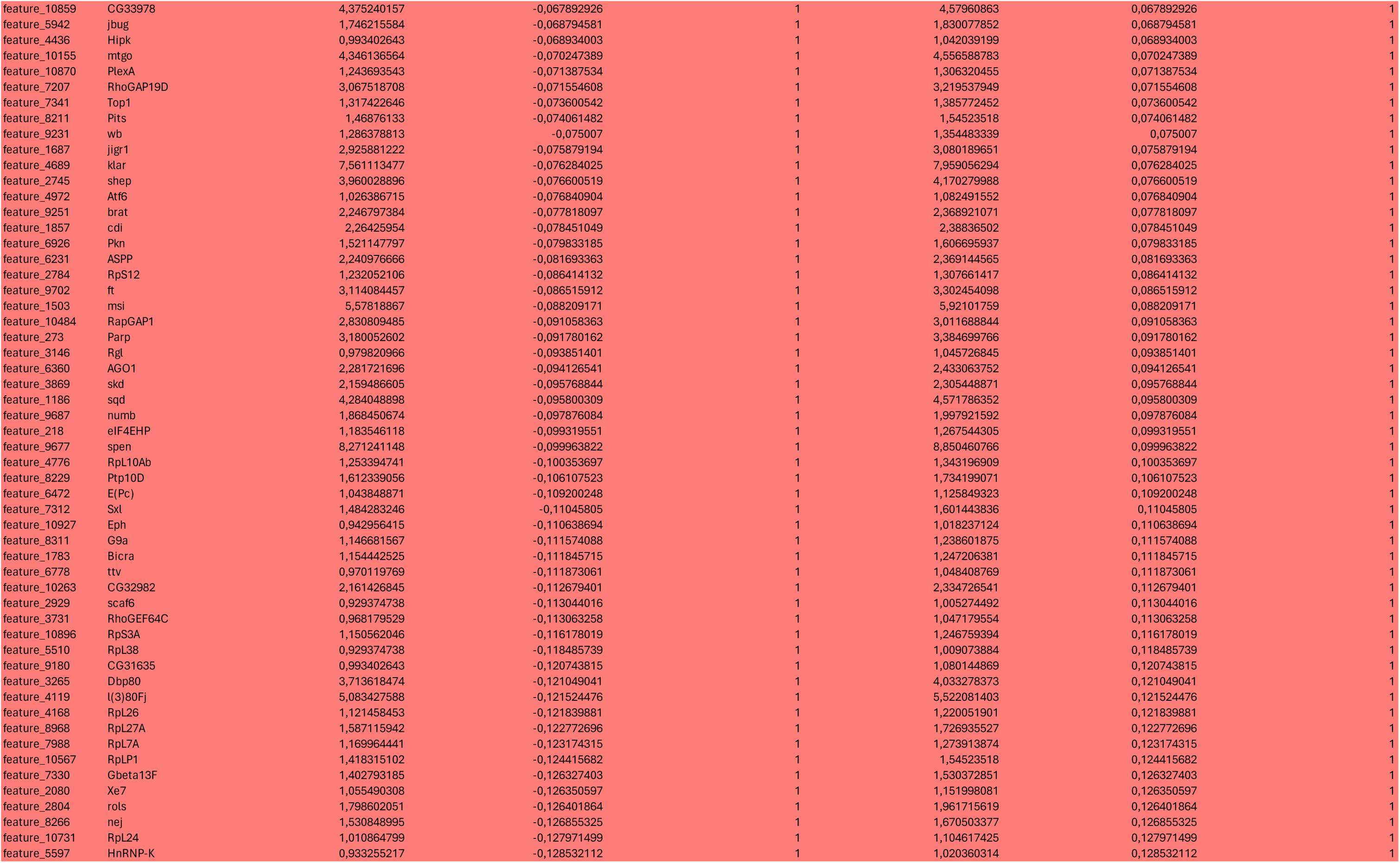
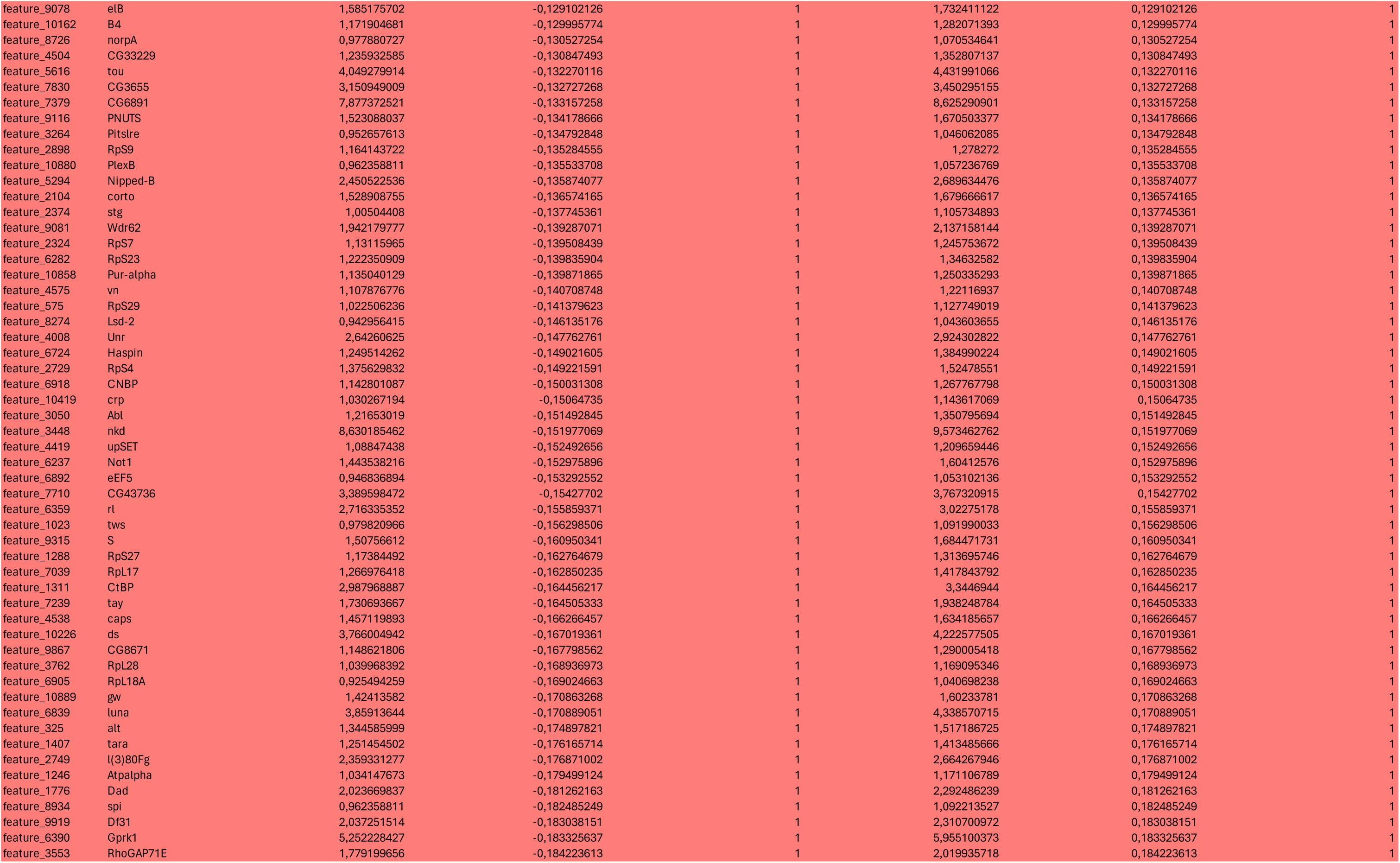
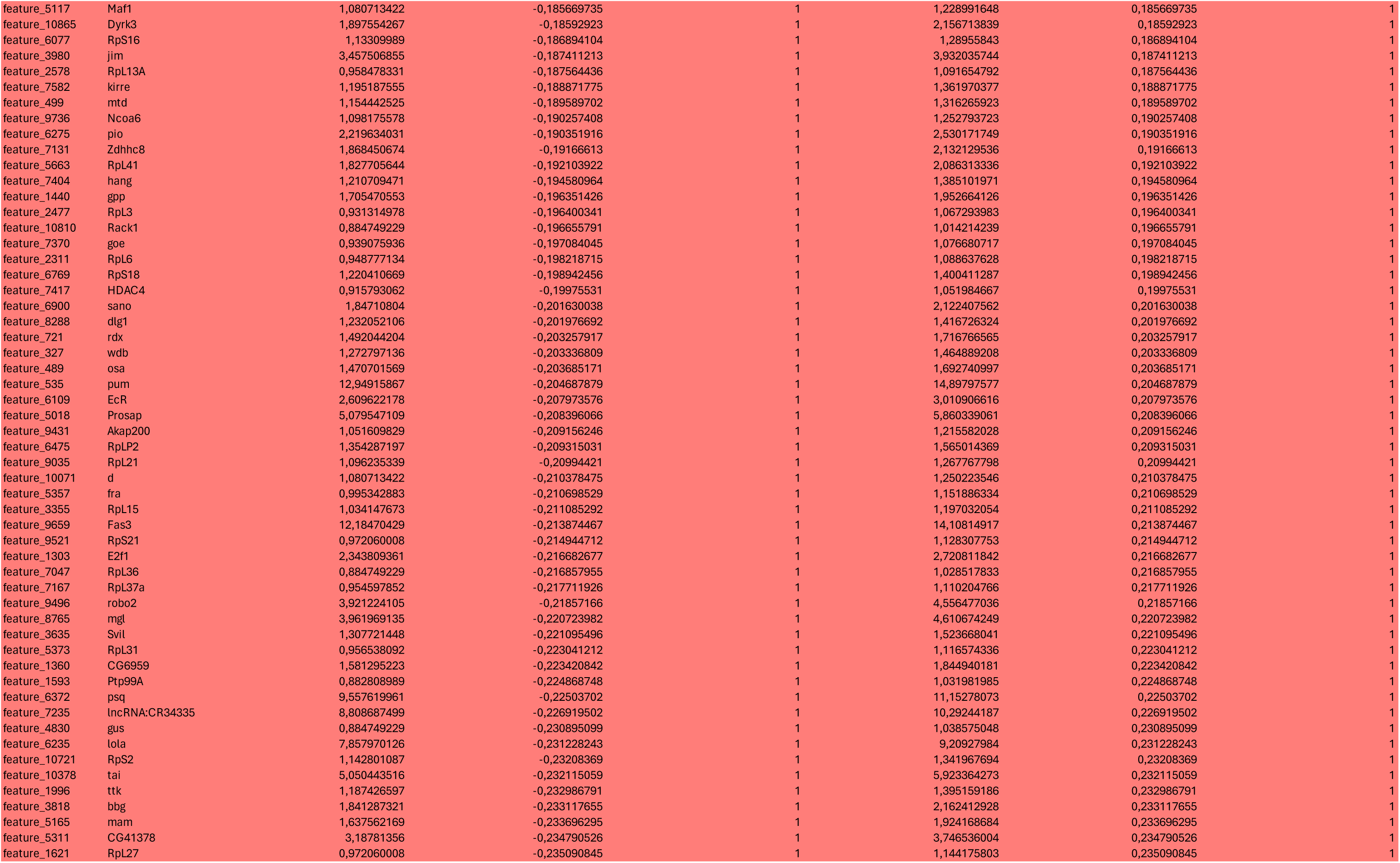
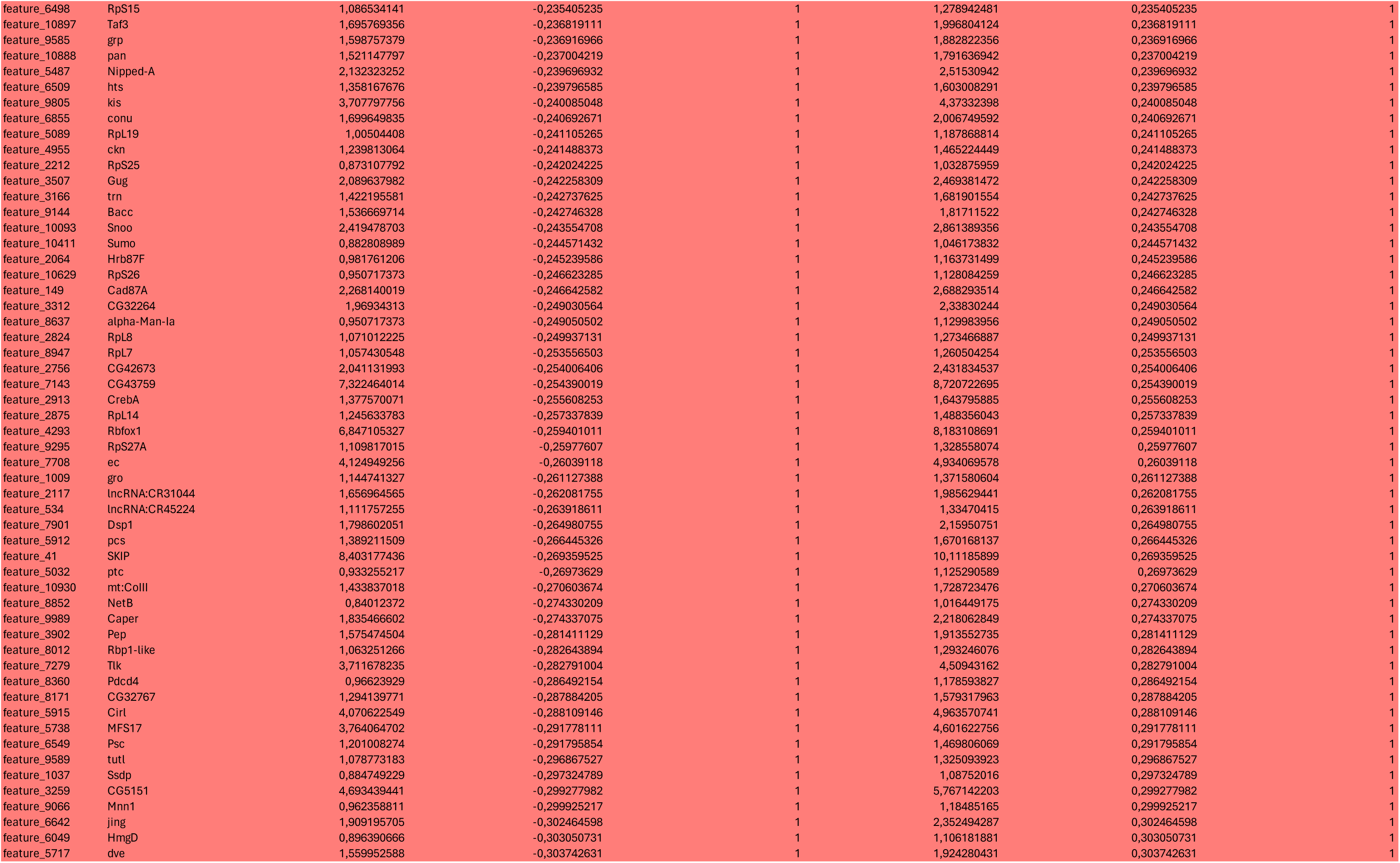
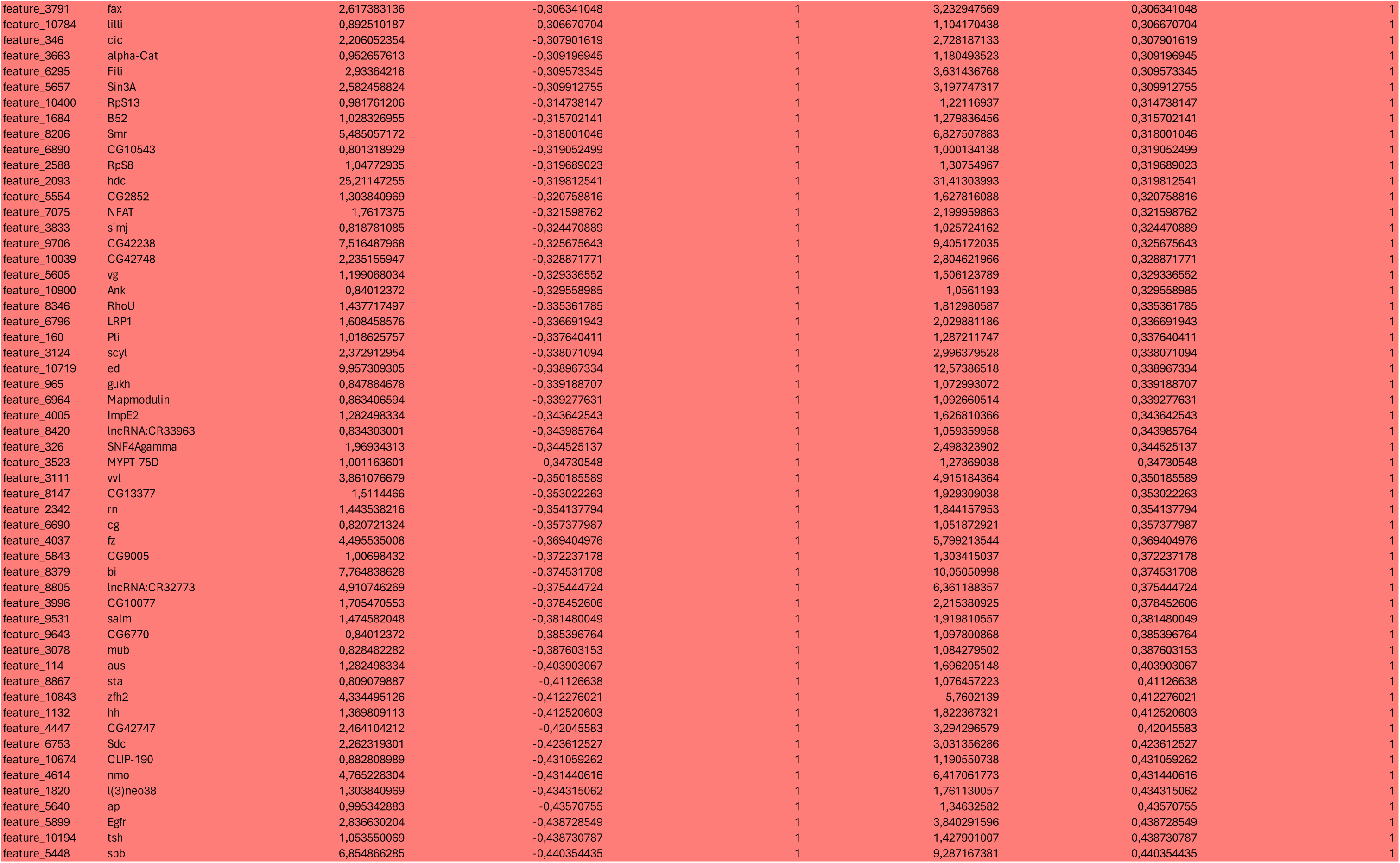
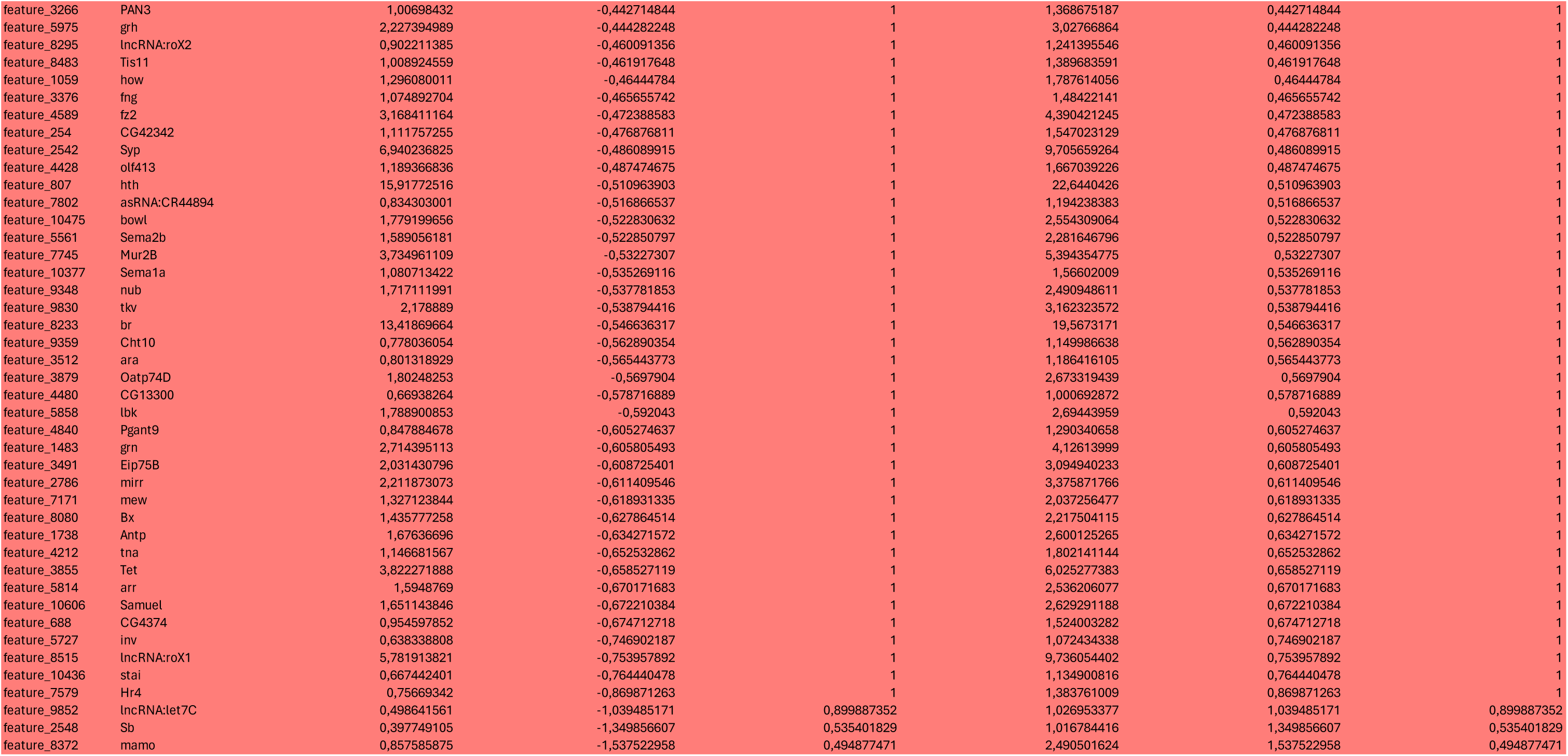
Differentially enriched or diminished expressed genes in the SPIKE subpopulation for Tumorigenic conditions.

Besides the establishment of a blastema (SPIKE) with tumorigenic character, repetitive local apoptosis results in a general derangement of the imaginal tissue. While a single apoptotic/regenerative event also stimulates blastema development, the contribution of the different wing subclusters to the global disc volume is proportionally unchanged. In the tumorigenic condition, however, the notum subclusters 1, 2 and 4 are strongly reduced, while the contribution of the Hinge-Outer 3 and PE 3 subclusters are enlarged (**Figure S4**). Thus, non-autonomous compensatory events emerge after successive rounds of aborted regeneration (e.g. anomalous growth rates outside the Rpr expressing domain) and contribute to the tumorigenic phenotype.

**Figure S2.**
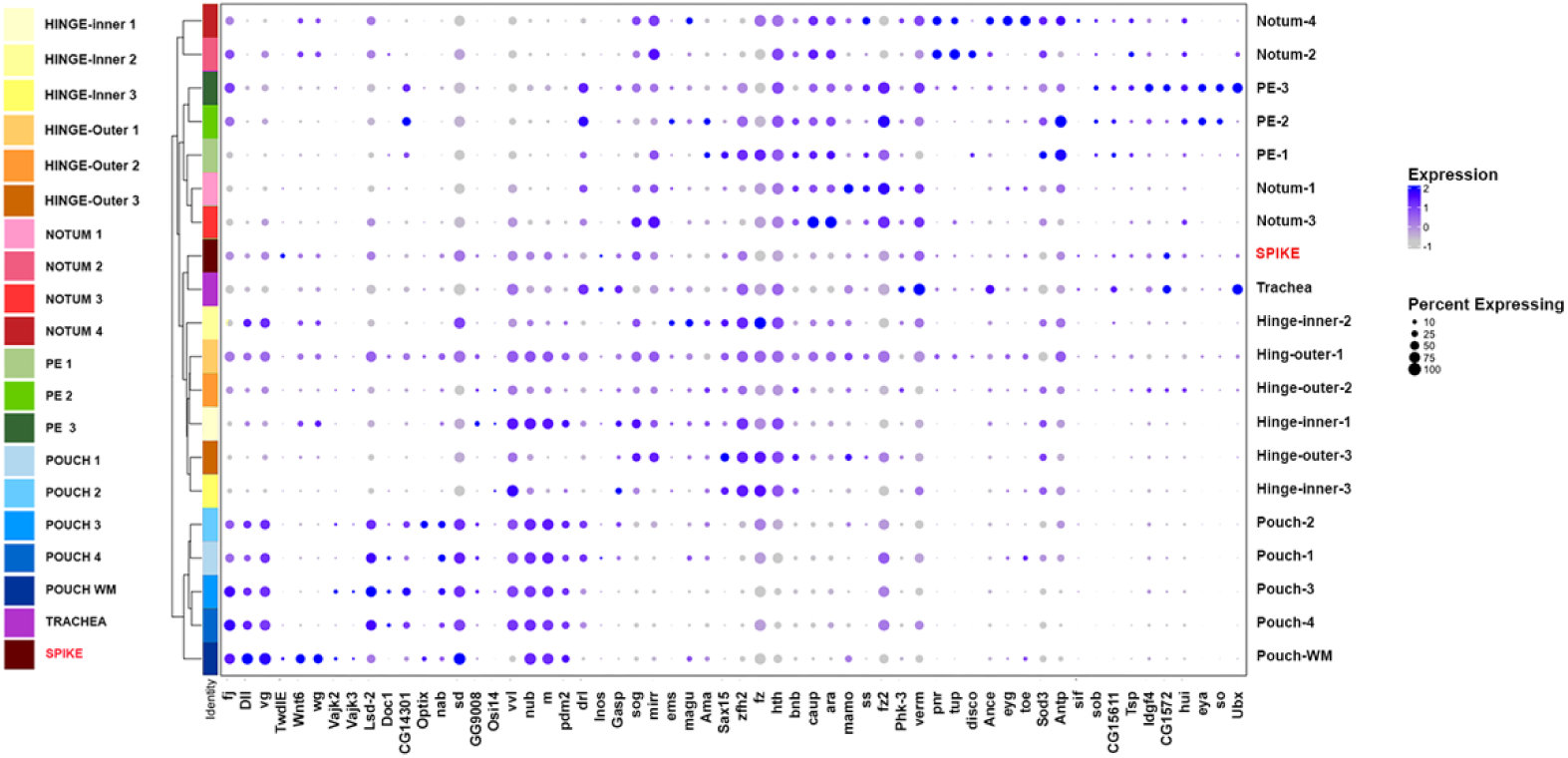
Patterns of expression of landmark genes. 58 landmark genes (expression level and expression percentage) define 30 subclusters within the overall snRNA Seq data. 19 of these clusters are represented (all except indirect and direct muscles subclusters). The new SPIKE subcluster is highlighted in red.

**Figure S3.**
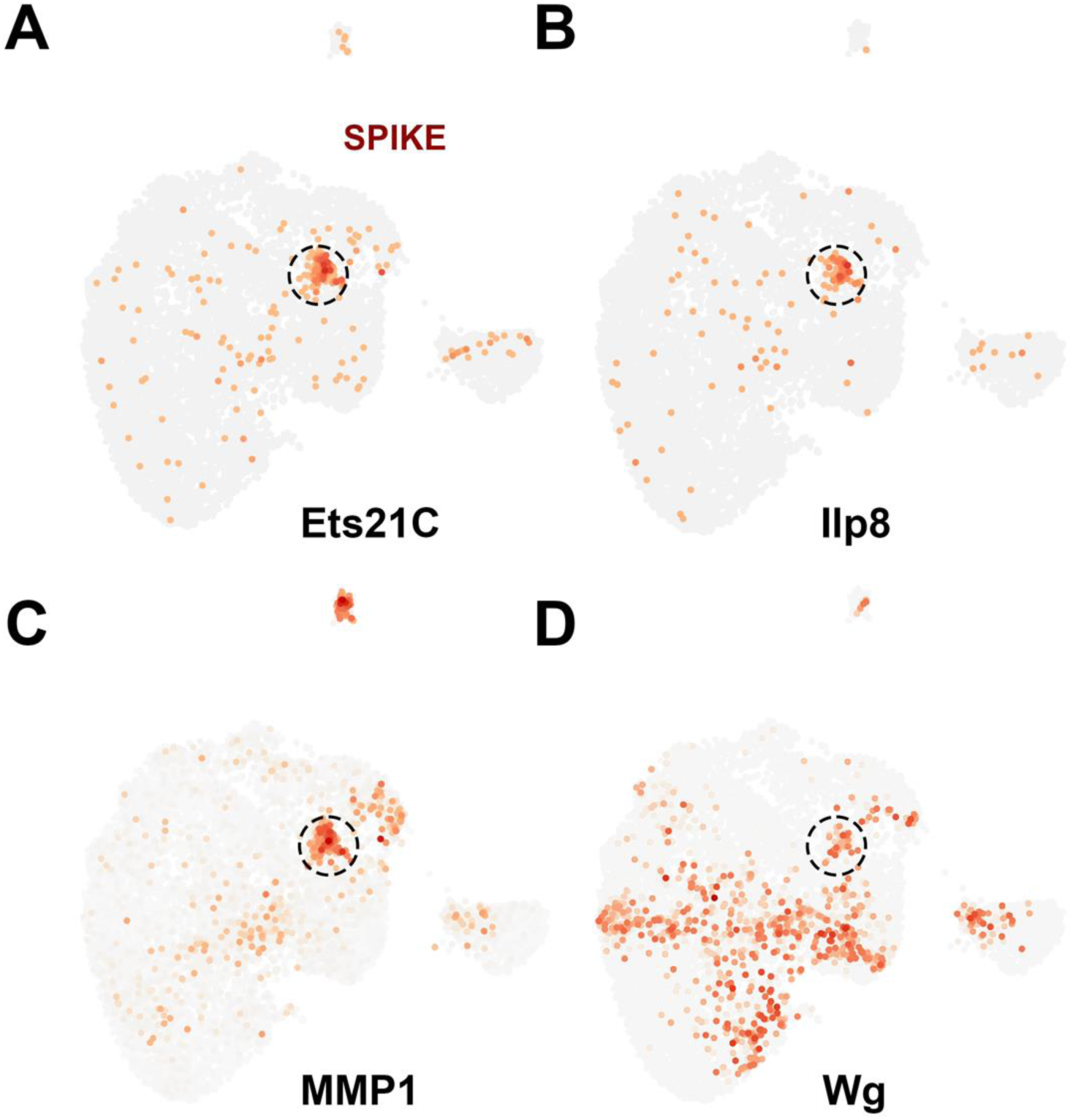
Feature plots highlighting expression of blastema enriched genes. **A)** Ets21C. **B)** Ilp8. **C)** MMP1. **D)** Wg. The SPIKE domain is encircled by a discontinued circumference.

**Figure S4.**
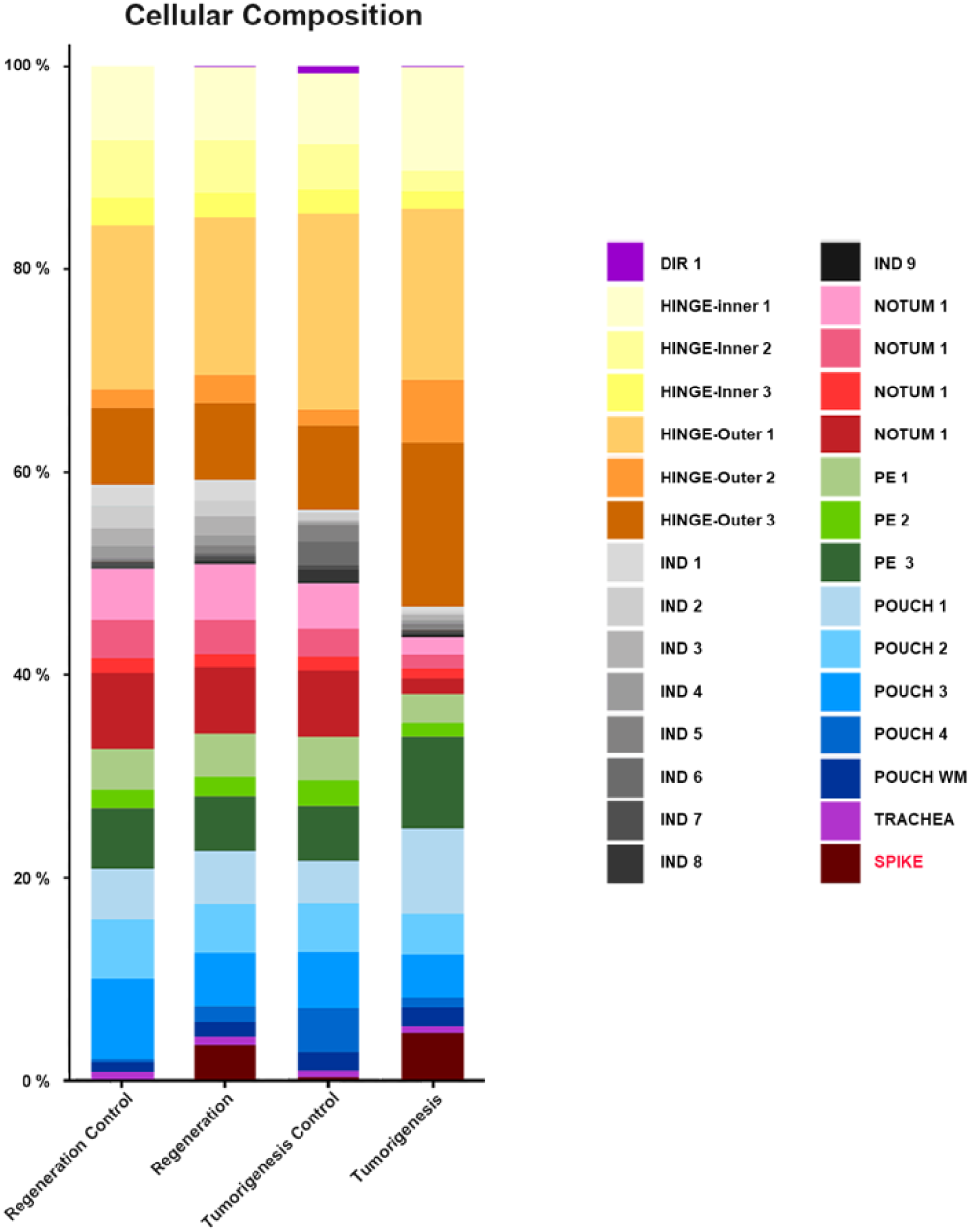
Percent subclusters composition of wing imaginal disc. Color coded marking of the different subcluster identified in the snRNAseq analysis for regeneration control, regeneration, tumorigenesis control and tumorigenesis wing discs. The SPIKE subcluster is shown in red.

### Role of JNK signaling in tumorigenic transformation

The SPIKE blastema transcriptional signature suggests that the JNK pathway constitutes a major element mediating tissue repair. Indeed, previous studies have established JNK in wing imaginal discs as a principal determining factor for normal regeneration upon induction of cell death ^16^. The Jun N-terminal kinase (JNK) belongs to the well-known mitogen-activated protein kinase (MAPK) family and in *Drosophila* is encoded by the gene *basket* (*bsk*). Bsk activity is modulated through a negative feedback loop mediated by *puckered* (*puc*), a gene coding for a dual-specificity phosphatase ^53^. JNK signaling display dual roles. On one hand, it activates the pro-apoptotic genes Rpr and Hid, which block the activity of the *Drosophila* inhibitor of apoptosis (Diap1), allowing the activation of the apical caspase Dronc. This leads to the subsequent activation of effector caspases, specifically Drice and Dcp1, causing the death of JNK-expressing cells. Dronc, in turn, stimulates the JNK activity, generating an amplification loop ^54^. On the other hand, JNK-expressing cells emit proliferative signals stimulating cell proliferation ^55^.

To evaluate how the JNK pathway may intervene in the neoplastic transformation of the regenerative blastema, we first monitored the level of JNK activity in the Ptc domain in discs subjected to different cycling death and regeneration regimes. To do so, we employed a genetically encoded biosensor (KTR-Jun), which identified those cells where the JNK cascade was active ^56^. Phosphorylation by JNK leads to the masking of an engineered nuclear localization signal and as a result the GFP-based sensor translocates from the nuclei to the cytoplasm. Expressing it in the death-targeted cells revealed, when compared to regenerative conditions, a strong activation of the pathway in the tumorigenic blastema (compare **Figures 8A** and **8B**). Following, we repressed JNK activity by co-expressing, along Rpr, a Bsk RNAi transgene . The overexpression of Bsk RNAi alone (controls) did not affect the growth or shape of the wing disc, neither the proliferation levels or number of DSBs (**Figure 8C**, **8E**, **8F** and **8H**). Yet, when co-expressed with Rpr, while the penetrance of the overgrown phenotype of Rpr-expressing discs was not affected, the presence of Bsk RNAi reduced the expansion of the Ptc domain and, strongly, mitosis numbers (**Figure 8G** and **8E**). Further, the number of DSBs suffered a significant increase (**Figure 8G** and **8H**). Thus, the activity of the JNK cascade after three cell death/regeneration cycles appears to boost proliferation and the tumorigenic transformation of the regenerative blastema. Also, it seems an important factor on assuring the efficiency of DSBs repair. While these two outcomes can look contradictory, an intensified response to a lack of repair competencies may prompt damaged cells to rapidly apoptose resulting on a diminished proliferating response (see Discussion).

### Tumorigenesis rescue by interference of the DNA repair machinery

The number of cells with double stranded DNA breaks (DSBs) detected after three rounds of induced cell death and regeneration increased significantly. This suggests a possible fundamental role of the DNA damage response (DDR) on neoplastic transformation of the regenerating blastema. To study if the alteration in the DNA repair system was causal for the transformed phenotype, we knocked down the expression of the *Drosophila* homologue (*tefu)* ^57^ of the ataxia-telangiectasia–mutated (ATM) gene. ATM is a central kinase that participate in the cellular response to DNA damage. ATM responds specifically to DSBs ^58^.

Larvae carrying Rpr and RNAi constructs for ATM (or these ones alone - control discs) were subjected to three cycles of apoptosis and regeneration as described. ATM RNAi controls were essentially normal. In these conditions, the shape of the discs and number of cells with DSBs were very similar to those of wild type siblings (**Figure 9A**, **9D** and **9F**). Surprisingly the reduction of ATM expression led to diminishing proliferation in ATM RNAi only controls (**Figure 9C**).

**Figure 9.**
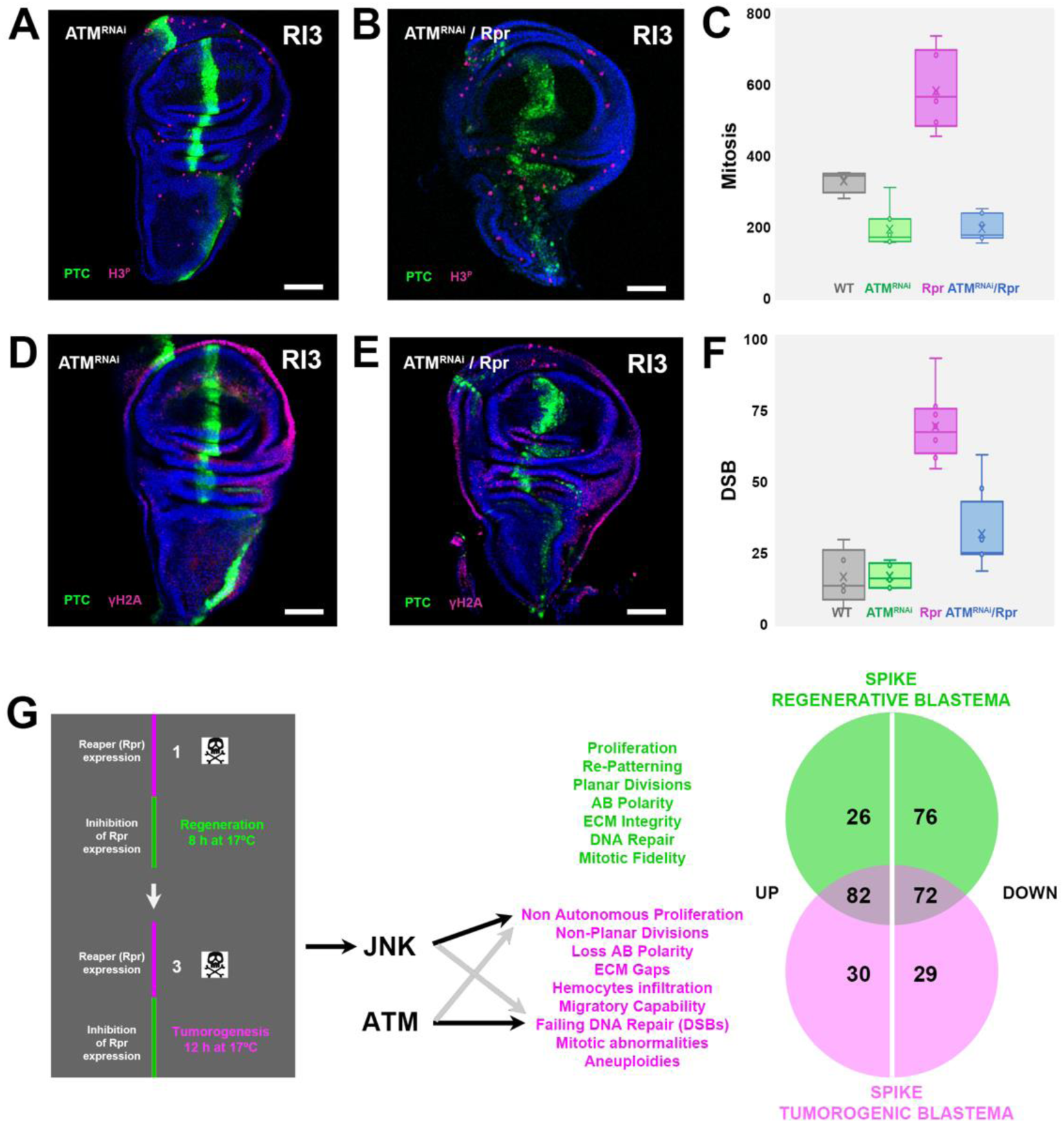
DNA repair and JNK signaling coordinately lead to blastema tumorigenic transformation. **A** and **B)** H3^P^ labelling (magenta) of RI3 wing discs depicting cells in mitosis after expressing an ATM^RNAi^ transgene (**A**) or after dual overexpression of ATM^RNAi^ and Rpr (**B**). Green is Ptc / GFP and blue is DAPI. Scale bar is 50 µm. **C)** Quantification of mitotic cells in wild type (grey) (n = 4), RI3 / ATM^RNAi^ (green) (n = 8), RI3 / Rpr (magenta) (n = 7) and RI3 / ATM^RNAi^ + Rpr (blue) (n = 7) wing discs. The two-tailed P value between RI3 / Rpr and RI3 / ATM^RNAi^ + Rpr is less than 0.0001, which is extremely statistically significant. **D** and **E)** γ-H2a labelling (magenta) of RI3 wing discs depicting cells with DSBs after expressing an ATM^RNAi^ transgene (**D**) or after dual overexpression of ATM^RNAi^ and Rpr (**E**). Green is Ptc / GFP and blue is DAPI. Scale bar is 50 µm. **F)** Quantification of cells presenting DSBs in wild type (grey) (n = 4), RI3 / ATM^RNAi^ (green) (n = 8), RI3 / Rpr (magenta) (n = 7) and RI3 / ATM^RNAi^ + Rpr (blue) (n = 8) wing discs. The two-tailed P value between RI3 / Rpr and RI3 / ATM^RNAi^ + Rpr equals 0.0006, which is extremely statistically significant. **G)** A model for the transformation of a regenerative blastema. Repetitive sequential induction of apoptosis at the anteroposterior compartment border in wing imaginal disc (RI3) leads to the hyperactivation of the JNK signaling cascade and to failures in DNA repair. This effectively transforms a natural regenerating blastema (green text) inducing tumorigenic characteristics (magenta text). This transformation associates to differential transcriptomic patterns of the blastemas (spike) with distinct upregulated and downregulated genes in regenerative and tumorigenic conditions.

Remarkably, the co-expression of Rpr and ATM RNAi employing equivalent protocols resulted in a partial rescue of both, the Rpr-induced tissue overgrowths and the expansion of the Ptc domain (**Figures 9B** and **9D**). It also leads to a strong reduction of ectopic mitosis to ATM RNAi discs levels, and to a decrease in the number of cells with DSBs, when compared to Rpr alone siblings (**Figure 9C** and **9F**). These results were puzzling as we were expecting an opposite outcome: i.e., that diminishing the ability to repair DSBs, by interfering into ATM expression, would increase DSBs and tumorigenic penetrance in cells challenged by Rpr. Still, these results could be explained as an exacerbated response to the lack of repair capabilities prompting damaged cells to rapid apoptosis (see Discussion). Anyhow, we can infer from these results that DNA damage control (DSBs) is an intrinsic element in the preservation, under repetitive stress, of the regeneration program.

## DISCUSSION

The first report pointing to a close relationship between healing and stress responses and tumors dates from 1863 ^59^. Rudolf Virchow, the founder of modern pathology, noted leukocytes in neoplastic tissues, and made a connection between inflammation and cancer suggesting that the “lymphoreticular infiltrate” reflected the origin of cancer at sites of chronic irritation. In the dawn of the 20th century, early studies on the pathogenesis of gastric carcinoma also led to suggest that chronic gastric ulcers may have a major role in the development of stomach cancer (irritation theory of cancer) ^59–61^. Since these early studies, a great number of related cases have been reported including metastases occurrence at sites of accidental trauma ^62^. In the face of these classic problem, we still are unaware of the cellular and molecular mechanisms affected when healing and regeneration fail leading to tumorigenesis. No detailed experimental analysis of the association between repair and regeneration and the origin of tumors has been reported so far.

Most of the signaling pathways controlling cell growth and migration in mammals have a conserved function in flies and this has allowed to develop several models in *Drosophila* that mimic tumor’s biology ^63^. These tumors have a growth rate much higher than surrounding wild type cells and display metastatic properties as appreciated when injected into adult flies.

*Drosophila* has lately reappeared as a good model for regeneration ^64^. The genetically controlled ablation of specific cellular domains within *Drosophila* imaginal discs stimulates the discs cells to change fates and fulfill complete tissue regeneration ^65^. We realized that the cellular plasticity of the regenerative blastema could be a good target for subverting the repair response of imaginal discs to go awry leading to tumor generation. Being this the case, this model would let to analyze “in vivo”, in an unprecedented way, the dichotomy between tissue repair/regeneration and tumor formation in response to injury. Our results showed that three sequential episodes of death induction/regeneration led to tissue neoplastic overgrowths that were not restricted to the apoptotic targeted area. Flies repeatedly challenged were unable to develop into normal adults and all died at pupal stages. The imaginal discs overgrowths generated as a failure of the regeneration process resulted from unpattern proliferation as the wing imaginal disc were unable to control mitosis rates. Cell overproliferation was detected, not just in or around the apoptotic targeted domain, but non-autonomously far away throughout the wing pouch, particularly on the anterior compartment, well known to respond to cell death by compensatory proliferation ^16^. Compensatory proliferation, a JNK-driven regenerative mechanism in which apoptotic or stressed cells induce mitogen release (Wg, Dpp, Upd, EGFR ligands) to stimulate neighboring cells to divide and restore tissue integrity ^66^. If excessive or chronic (e.g. persistent “undead” cells), it can drive hyperplasia and tumor-like growth ^67^. Hence, it represents a physiological regenerative mechanism that can become pathological if unchecked (e.g. when apoptosis is blocked or JNK remains active).

We further characterized these neoplastic overgrowths and found that they are composed of cells with interstitial migratory capacity originating within the regenerative competent domain, including undead cells from the apoptosis targeted area (“walking dead cells”) (**Figure 9G**). In some way, these tumors were similar to those generated upon X-ray irradiation in conditions in which cell death was abolished ^68^. However, our tumorigenic model will not demand to let cells to survive, it is the consequence of a natural response of the tissue to repeated injuries.

In general, tumors generate as a result of the deregulation of protooncogenes and the evasion of the anti-proliferative effects of tumor suppressors controlling cell cycle check points. Cancer cells proliferate and replicate unlimitedly and avoid apoptotic death, eluding cellular homeostasis, by increasing the activity of anti-apoptotic genes and pro-survival factors and/or by downregulating the action of pro-apoptotic genes. They become invasive acquiring metastatic capabilities by losing their apical-basal cell polarity along the degradation of the extracellular matrix (ECM) in a process altering the adjacent microenvironment, which may provide non-autonomous signals supporting growth ^69^. While not a formal proof for metastasis, the presence of “walking dead cells” upon recurrent death induction indicates that many of them actively intercalate throughout the epithelia and may have metastatic potential.

Which are the causal mechanisms promoting the aberrant behavior of the regenerating blastema? If the observed neoplastic overgrowths resulted from a failure to apoptose of those cells with an aberrant DNA content, we would expect that manipulating the DNA repair machinery will affect the penetrance and/or the expressivity of these phenotypes. We found that interfering with the DNA repair response or the activity of the JNK signaling in the targeted tissue leads to drops on both, the tumorigenic penetrance and uncontrolled proliferation (**Figure 9G**).

The DNA damage response (DDR) is essential for proper cell divisions and rescues cells from aberrant detours in their physiology, preventing genome instability ^70^. DSBs, which are the main lesions that the DNA suffers during replication, are recognized by sensors (MRN complex) that recruit the DDR proteins, including the ATM kinase (Tefu in *Drosophila*), to the damaged site. ATM becomes activated and promotes repair by phosphorylation and recruitment of other substrates including the checkpoint kinase CHK2. This leads to an arrest of the cell cycle and prompts to error-free homologous recombination (HR) ^71^. When DNA damage is overly strong, the DDR opts, alternatively, to induce senescence or apoptosis ^72,73^. In flies, upon low-dose irradiation, the Tefu–Chk2 partnership is mainly responsible for p53-dependent apoptosis and can induce cell cycle arrest ^74–76^. When apoptosis is suppressed, irradiation of wing imaginal discs causes tumorigenic growth ^77^. In this context, the partial suppression of repeated cell death-induced tumorigenesis by interfering with ATM expression could reflect an increment in apoptosis for those cells engaged in neoplastic transformation, further, decreasing the ability to repair DSBs and leading to a reduction of proliferation rates and the number of aberrant “walking dead cells”.

The role of JNK in tumorigenesis in *Drosophila* is dithering, and fulfills different, at first glance, opposing roles. JNK signaling participates in the elimination of pre-tumorigenic cells via apoptosis, but also it can promote neoplastic transformation ^78,79^. JNK signaling can be activated by the Tumor Necrosis Factor (TNF), or after cellular damage inflicted by reactive oxygen species (ROS), promoting the caspase-mediated death of tumorigenic cells. It also participates in the elimination, by means of cell competition, of mutant cells for polarity genes such as *scribble*, which can promote neoplastic transformations.

In our tumorigenic model, inhibiting JNK deters tumor growth, but how this works? The Hippo/Yorkie (Yki) pathway is an important growth regulator, which modulates targets such as the cell cycle regulator CycE or the apoptosis inhibitor Death associated inhibitor of apoptosis 1 (Diap1) ^80^. Induction of apoptosis in the *Drosophila* wing disc, or the depletion of neoplastic tumor suppressor polarity genes, stimulates the activation of Yki, which is required for regeneration, via JNK ^81^. Blocking JNK signaling results in Yki inhibition and upregulation of Yki targets ^82^. One appealing possibility is that the sequential activation of programmed cell death will lead to the unbalanced activity of Yki in response to spaced pulses of JNK activity. A potential different life-expectancy of Yki targets, promoting either cell divisions or apoptosis inhibition, may lead to overproliferation and apoptosis resistance, and ultimately neoplastic overgrowths. Blocking JNK expression could then moderate these after-effects. A further indication pointing to Yki as a potential mediator of the aberrant response of wing imaginal cells to sequential death inputs, is its ability to bypass JNK-mediated tumorigenesis prevention in response to cytokinesis failure and aneuploidies (a condition shown to upregulate JNK signaling) ^83^. Last, hemocyte attraction has been shown to depend on JNK-mediated secretion of a cleaved form of the protein Tyrosyl-tRNA synthetase ^84^. If this would be the case in our experimental conditions, inhibiting JNK expression would lead, as found, to tumorigenic relapse.

We do not know if the JNK pathway and the DNA repair response are somehow interconnected. Tumorigenesis in imaginal discs is triggered by JNK signaling upon irradiation. DDR, in this context, functions as a tumor suppressor mechanism and HR DNA repair and cell cycle arrest are crucial for tumorigenic suppression ^85^. Further, the inhibition of the G2/M regulator cdc25/string, a target of JNK, suppresses tumorigenic growth ^70^. The DDR and JNK signaling could thus be interlinked and determine together whether a cell undergoes cell cycle arrest triggering HR DNA repair or it is terminally damaged and either undergoes apoptosis or get engaged in neoplastic overgrowths.

Our snRNA-seq analysis resolves a discrete, stress-responsive SPIKE population that is absent in controls and emerges after repeated injury or tumor induction. In both tumorigenesis and regeneration conditions, SPIKE nuclei occupy an identical position between NOTUM and PE in UMAP space and share a wound/regeneration signature (e.g., Ilp8, Mmp1, Wg, Ets21C) consistent with a damage-licensed state at the repair front reminiscent of previously described blastema-like cells ^52^. Further, the JNK signaling cascade comes out as a mark of SPIKE. Beyond this shared identity, SPIKE bifurcates transcriptionally by condition: regenerative SPIKE is biased toward heat-shock programs, whereas tumorigenic SPIKE is enriched for specific JNK pathway components, aligning with our functional data showing that JNK activity promotes overgrowth. Together, these findings support a model in which repeated injury stabilizes a JNK-high SPIKE state with expanded proliferative/migratory potential, predicting that targeting JNK signaling or the DNA-damage response will subside SPIKE and restore regenerative restraint.

## EXPERIMENTAL PROCEDURES

### Apoptosis cycle protocol: steps, immunostaining and dissection

Crosses were made between appropriate genetically designed stocks (see below). The selected descendants expressed GFP and Rpr in the *ptc* pattern and carry a thermosensitive allele of the Gal4 repressor, Gal80^ts^. 4 h embryo collections on agar from flies kept at 25°C were transferred to 17° C to block continuous Rpr expression. One day later (24 h) the agar piece containing eggs was transferred into a vial with fly food and kept at 17° C during five more days, when the larvae reach their second stage. At this time, the vials were relocated to 29° C for 16 h to induce Rpr expression and apoptosis, inhibiting the Gal80^ts^. Transferring the tubes back to 17° C stopped apoptosis induction and the larvae were allowed to recover during 8 h. A second 29° C period of 16 h was set. Recovery at 17° C for 8 h was again performed and a third round of incubation at 29°C executed for 16 h. In those cases, in which tissue recovery after three rounds of apoptosis was evaluated, the larvae were kept at 17° C during 12 extra hours before dissection. In flies undergoing a single apoptosis/regeneration round, the transfer to 29° C was performed once the animals reached the third instar larval stage after 8 days at 17° C.

Larvae of the appropriate genotypes were selected and put in a dissection plate with PBS 1X. One by one, larvae were dissected and the wing imaginal discs, attached to the larval carcass, collected. The PBS 1X medium was exchanged to Formaldehyde 4 % in PBS 1X and an incubation with gentle shaking was performed for 30 minutes at room temperature (RT) to fix tissues. Medium was exchanged to PBS-Tween (0.1 %) (PBS-T) to permeabilize the tissue, four times 15 minute each, at RT. The fixed larvae blocked with Bovine Serum Albumin (BSA) 5 % in PBS-T during one hour at RT and first antibodies were added (diluted in PBS-T BSA). The sample was kept at 4° C overnight. Next day, the carcasses were washed four times with PBS-T (15 minutes each) and then the second conjugated antibody diluted in PBS-T was added and kept during 1 h and 30 min at RT. Samples were washed two times with PBS-T and stained with DAPI (1:1000 in PBS-T) during 15 min at RT without shaking. DAPI was cleaned away, washing once with PBS-T and once with PBS 1X. The carcasses were then carefully dissected in a plate with PBS 1X and the wing discs isolated. Discs were pipetted using non sticky tips into a cover plate previously treated with polylisine. When wing discs got attached to it, the PBS 1X was removed and the discs were mounted on a drop of Vectashield.

### G-TRACE

The Gal4 technique for real-time and clonal expression known as G-TRACE [13] was used to study lineage cell tracking. It combines three elements: a UAS-RFP fluorescent protein (red), a UAS-flipase and a Ubi-FRT-STOP-FRT-GFP construct (green) recombined in a single chromosome. We generated a stock carrying a UAS-Rpr construct in the X chromosome and a G-TRACE combination in the 2nd chromosome. Females of this stock were crossed with males from the Gal4 stock under the control of Ptc recombined with Ubi-Gal80ts. From this cross the expression of the UAS-Flipase (Flp) will initiate in the progeny as soon as the Gal4 expression started. Thus, from the beginning, the FLP will induce the excision of the FRT cassette and all cells once expressing Ptc will be permanently labelled in green. The red labelling of the ptc- Gal4 expressing cells induced from the UAS-RFP reporter will fade away over time and will only be detected in those cells expressing the Gal4 at the time of the analysis or shortly before. For these analyses, selected larvae were fixed, permeabilized and stained with DAPI before dissecting the wing discs.

### Disaggregating wing disc cells and detection of aneuploids by FACS

Control (no Rpr) and experimental (Rpr) 3 RT periods + 12 h) larvae from the apoptosis/regeneration crosses were selected and their wing imaginal discs carefully dissected in PBS 1X. About 50-60 wing discs (manually selected overgrown ones in the case of the 3 periods + 12 h larvae) were placed in an Eppendorf tube with 400 μL of PBS 1X. Then 100 μL of Trypsin-EDTA 10X were added and the Eppendorf incubated at 32° C during 45 minutes to activate trypsin. The sample was passed through a 23G needle twice; after 20 and 45 min of trypsinization to facilitate disaggregation. Then 60 μL of fetal bovine serum (FBS) were added to stop trypsin activity and 400 μL of PBS 1X and 50 μL of formaldehyde, afterwards, to fix the sample for 20 minutes shaking at RT. Cells were centrifuged 5 min at 7000 rpm and the supernatant decanted carefully. Cells in the pellet were resuspended in 100 μL of PBS 1X and then 1 mL of ethanol 70 % was added drop by drop. This mix was kept for 1 h without shaking at RT. Cells were centrifuged again with the same parameters and resuspended in 500 μL of DAPI solution (triton X-100 0.1 % in PBS 1X + 0,5 μL DAPI 1 mg/mL + 2 μL Hoesch 1 mg/ml + 1 μL RNAse A 100 mg/mL [final]=0.2 mg/ml). The sample was kept 4° C overnight without shaking.

In the sorter we distinguished cells that expressed GFP (from the *ptc* domain of the wing disc) and determined their DNA content (by the level of DAPI signal). Cells that are in G1 phase or in G2 phase can be easily recognized (double amount of DNA, double amount of DAPI signal). Cells with carry more than 2 copies of the DNA (aneuploids) were quantitatively assessed.

### Larvae dissection and sample preparation

Following the rounds of apoptosis, dissection of the wing imaginal discs was performed on third instar larvae of control and experimental flies. Dissected discs were transferred to 100 μl of medium in a nuclease-free 1.5 ml Eppendorf tube kept on ice. Approximately 100 discs were collected per sample. After dissection, samples were spin down, the medium was replaced with 10 μl of PBS 1X, and samples were flash-frozen in liquid nitrogen and stored at –80 °C.

### Single-nuclei RNA sequencing

Flash-frozen fly wing imaginal disc samples (50 each) were homogenized in a 1 ml Dounce in buffer consisting of 250 mM sucrose,10 mM Tris pH 8.0, 25 mM KCl, 5 mM MgCl, 0.1 % Triton-X, 0.5 % RNasin plus (Promega, N2615), 1× protease inhibitor (Promega, G652A), 0.1 mM DTT, then filtered through 40-μm cell strainer and 40 μm Flowmi (BelArt, H13680-0040). Samples were centrifuged, washed and resuspended in PBS 1X with 0.5 % BSA and 0.5 % RNasin plus. The suspension was filtered again with 40 μm Flowmi immediately before FACS sorting. Nuclei were stained with DAPI Dye and sorted using BD FACSAria™ Flow Cytometer at the Flow Cytometry Facility at Harvard Medical School. After sorting, nuclei were collected and resuspended at 700–800 cells μl–1 in PBS 1X buffer with 0.5 % BSA and 0.5 % RNasin plus.

The snRNA-seq library was generated according to the 10X Genomics protocol (Chromium Next GEM Single Cell 3’_v3.1_Rev_D). Sequencing was conducted using Illumina NovaSeq at the Harvard Medical School Biopolymers Facility.

### Bioinformatic analysis

We processed the snRNA-seq data using Cell Ranger count pipeline version 8.0.1 using the *Drosophila melanogaster* reference genome BDGP 6.32. The feature-barcode matrices were further processed in R (version 4.3.2) using Seurat (version 5.0.2). Low-quality nuclei with a unique molecular identifier count of < 500 and mitoRatio > 10 % were filtered out and genes expressed in < 10 cells were removed. Ambient RNA correction was performed using SoupX (version 1.6.2). The data was then processed as previously described ^86^.

Marker gene identification was used to annotate 30 unique clusters in the dataset. Differential expression analysis was carried out for each cell type using a Wilcoxon rank-sum test between regenerative flies and non-regenerative flies for both tumor and control samples. All single-nuclei metrics were generated using ggplot2 (version 3.5.0).

## LEAD CONTACT

Requests for resources should be directed to, and will be fulfilled by, the Lead Contact, Enrique Martin-Blanco.

## MATERIALS AVAILABILITY

All fly strains originated from the Bloomington Drosophila Stock Center (BDSC), Kyoto Drosophila Stock Center (DGCR), Vienna Drosophila Resource Center (VDRC) or previously generated in the laboratory ^56^ (see Key Resources Table).

## DATA SHARING AND RESOURCE AVAILABILITY

Raw snRNA-seq reads have been deposited in the NCBI Gene Expression Omnibus (GEO) database (accession code will be available upon publication). Processed datasets can be mined through a web-tool [https://www.flyrnai.org/scRNA/] that allows users to explore genes and cell types of interest.

## ACKNOWLEDGMENTS

We would like to thank Florenci Serras (UB), Marco Milan (IRB), Monica Betancourt-Diaz (IGC) and Michel Boutros (DKFZ) for experimental suggestions and to members of the Martin-Blanco lab for thoughtful feedback and discussion on the manuscript. This work was supported by grants BFU2017-82876-P, PID2020-116273GB-I00 and PID2023-146038NB-I00 to EMB.

YL is supported by Charles A. King Trust. NP is an investigator of the Howard Hughes Medical Institute.

## AUTHOR CONTRIBUTIONS

Conceptualization, EMB; methodology, TK, AR, YL, PD, JU, LRE and KK; investigation, TK, AR, YL, PD, JU, LRE and KK; software, YL; formal analysis, EMB; visualization, TK, AR, YL; writing - original draft, EMB; writing - review and editing, EMB and NP; funding acquisition, EMB and NP; supervision, EMB.

## DECLARATION OF INTERESTS

The authors declare no competing interests.

This article is subject to HHMI’s Immediate Access to Research policy, which requires that this article be made publicly available as initial and revised preprints deposited on a designated preprint server under a CC BY 4.0 license.

## KEY RESOURCES

| Fly stocks | Source | Chromosome | Comments |
| --- | --- | --- | --- |
| w; Ptc-Gal4, UAS-GFP; tub-Gal80ts / SM5A TM6B | this paper | II,III | For regeneration/tumorigenic experiments |
| UAS-w1118 P{UAS-rpr.C}27 | BDSC 5823 | X | To monitor dlg expression |
| w1118 |  | X | Wild type Control |
| dlg-GFP; ptc-Gal4, UAS-RFP; tub-Gal80ts / SM5A TM6B | this paper | X,II,III | To monitor dlg expression |
| UAS-Rpr; DE-Cad::GFP / CyO, Dfd-YFP; tub-Gal80ts / SM5A TM6B | this paper | X,II | To monitor DE-Cad expression |
| UAS-Rpr; GTRACE / CyO | this paper | X,II | For GTRACE lineage |
| Ptc-Gal4 /CyO; UAS-JNK-KTR | this paper | II,III | To monitor JNK activity |
| UAS-Rpr; H2A RFP / If; Gal80ts / MKRS | this paper | X,II,III | To monitor JNK activity |
| UAS-bsk RNAi | VDRC (V34138) | II | To inhibit Bsk expression |
| UAS-Rpr / FM6; UAS-Bsk RNAi / (CyO | this paper | X,II | For Bsk RNAi tumorigenic test |
| UAS-ATM RNAi / (CyO) | VDRC (V22802) | II | To inhibit ATM expression |
| UAS-Rpr / FM6; UAS-ATM RNAi / (CyO) | this paper | X,II | For ATM RNAi tumorigenic test |
| y, w; hhPyR215 / TM3 (hh-dsRed) | DGCR (109137) | III | hh expression |
| UAS-Rpr;; hhPyR215 / TM3 | this paper | X, III | To monitor hh expression |

| Antibodies and immunocytochemistry Reagents | Source | Dilution | Comments |
| --- | --- | --- | --- |
| Mouse Anti-GFP | A-11120 (Invitrogen) | 1:600 | For Ptc expressing cells |
| Rabbit Anti-GFP | A-11122 (Invitrogen) | 1:600 | For Ptc expressing cells |
| Rabbit Anti-H3P | 9701s (Cell Signaling) | 1:600 | Mitosis |
| Mouse Anti-α-Tubulin-FITC | F2168 (SIGMA) | 1:200 | Spindles |
| Rabbit Anti-γ-Tubulin | T5192 (SIGMA) | 1:500 | Centrosomes |
| Rabbit Anti-Dcp-1 | 9578s (Cell Signaling) | 1:100 | Apoptotic cells |
| Mouse Anti-γ-H2A | UNC93-5.2.1-s (DSHB) | 1:100 | DNA damage |
| Mouse Anti-Crumbs | Cq4 (DSHB) | 1:100 | Crumbs |
| Mouse Anti-DE-Cadherin | DCAD2 (DSHB) | 1:20 | DE-Cadherin |
| Mouse Anti H2 (Hemese) | Dm 1.2/1 (DSHB) | 1:100 | Macrophages |
| ci | 2A1 (DSHB) | 1:100 | Ci |
| wg | 4D4 (DSHB) | 1:100 | Wg |
| TO-PRO 3 | R37170 (Thermo Fisher) | 1 μM | Nuclei |
| DAPI | D9542 (Sigma) | 300 nM | Nuclei |
| Goat Alexa fluor anti-Mouse 488 | A-11029 (Invitrogen) | 1:600 | Fluorescent Secondary Antibody |
| Goat Alexa fluor anti-Mouse 555 | A-21424 (Invitrogen) | 1:600 | Fluorescent Secondary Antibody |
| Goat Alexa fluor anti-Rabbit 488 | A-11034 (Invitrogen) | 1:600 | Fluorescent Secondary Antibody |
| Goat Alexa fluor anti-Rabbit 555 | A-21429 (Invitrogen) | 1:600 | Fluorescent Secondary Antibody |

## REFERENCES

1. Arwert, E.N., Hoste, E., and Watt, F.M. (2012). Epithelial stem cells, wound healing and cancer. Nat Rev Cancer 12, 170–180..

2. Feng, Y., Santoriello, C., Mione, M., Hurlstone, A., and Martin, P. (2010). Live imaging of innate immune cell sensing of transformed cells in zebrafish larvae: parallels between tumor initiation and wound inflammation. PLoS Biol 8, e1000562.

3. Friedl, P., and Gilmour, D. (2009). Collective cell migration in morphogenesis, regeneration and cancer. Nat Rev Mol Cell Biol 10, 445–457.

4. Schafer, M., and Werner, S. (2008). Cancer as an overhealing wound: an old hypothesis revisited. Nat Rev Mol Cell Biol 9, 628–638.

5. Coussens, L.M., and Werb, Z. (2002). Inflammation and cancer. Nature 420, 860–867.

6. Paro, R., Grossniklaus, U., Santoro, R., and Wutz, A. (2021). Regeneration and Reprogramming. In Introduction to Epigenetics, R. Paro, U. Grossniklaus, R. Santoro, and A. Wutz, eds. (Springer International Publishing), pp. 135–149..

7. Álvarez-Fernández, C., Tamirisa, S., Prada, F., Chernomoretz, A., Podhajcer, O., Blanco, E., and Martín-Blanco, E. (2015). Identification and functional analysis of healing regulators in Drosophila. PLoS genetics 11, e1004965.

8. Bosch, M., Serras, F., Martín-Blanco, E., and Baguñà, J. (2005). JNK signaling pathway required for wound healing in regenerating Drosophila wing imaginal discs. Developmental biology 280, 73–86.

9. Harris, R.E., Stinchfield, M.J., Nystrom, S.L., McKay, D.J., and Hariharan, I.K. (2020). Damage-responsive, maturity-silenced enhancers regulate multiple genes that direct regeneration in Drosophila. Elife 9, e58305.

10. Blanco, E., Ruiz-Romero, M., Beltran, S., Bosch, M., Punset, A., Serras, F., and Corominas, M. (2010). Gene expression following induction of regeneration in Drosophila wing imaginal discs. Expression profile of regenerating wing discs. BMC Developmental Biology 10, 94.

11. Repiso, A., Bergantinos, C., Corominas, M., and Serras, F. (2011). Tissue repair and regeneration in Drosophila imaginal discs. Dev Growth Differ 53, 177–185.

12. Igaki, T., Pagliarini, R.A., and Xu, T. (2006). Loss of Cell Polarity Drives Tumor Growth and Invasion through JNK Activation in Drosophila. Current Biology 16, 1139–1146.

13. Enomoto, M., and Igaki, T. (2011). Deciphering tumor-suppressor signaling in flies: Genetic link between Scribble/Dlg/Lgl and the Hippo pathways. Journal of Genetics and Genomics 38, 461–470.

14. Miles, W.O., Dyson, N.J., and Walker, J.A. (2011). Modeling tumor invasion and metastasis in Drosophila. Dis Model Mech 4, 753–761.

15. Kumar, R., and Hong, W. (2024). Hippo Signaling at the Hallmarks of Cancer and Drug Resistance. Cells 13, 564.

16. Bergantinos, C., Corominas, M., and Serras, F. (2010). Cell death-induced regeneration in wing imaginal discs requires JNK signalling. Development 137, 1169–1179.

17. Repiso, A., Bergantinos, C., and Serras, F. (2013). Cell fate respecification and cell division orientation drive intercalary regeneration in Drosophila wing discs. Development 140, 3541–3551.

18. Lu, B., Roegiers, F., Jan, L.Y., and Jan, Y.N. (2001). Adherens junctions inhibit asymmetric division in the Drosophila epithelium. Nature 409, 522–525.

19. Gibson, M.C., Patel, A.B., Nagpal, R., and Perrimon, N. (2006). The emergence of geometric order in proliferating metazoan epithelia. Nature 442, 1038–1041.

20. Reinsch, S., and Karsenti, E. (1994). Orientation of spindle axis and distribution of plasma membrane proteins during cell division in polarized MDCKII cells. J Cell Biol 126, 1509–1526.

21. Morin, X., and Bellaiche, Y. (2011). Mitotic spindle orientation in asymmetric and symmetric cell divisions during animal development. Dev Cell 21, 102–119.

22. Nakajima, Y., Meyer, E.J., Kroesen, A., McKinney, S.A., and Gibson, M.C. (2013). Epithelial junctions maintain tissue architecture by directing planar spindle orientation. Nature 500, 359–362.

23. Fleming, E.S., Temchin, M., Wu, Q., Maggio-Price, L., and Tirnauer, J.S. (2009). Spindle misorientation in tumors from APC(min/+) mice. Mol Carcinog 48, 592–598.

24. Vasiliev, J.M., Omelchenko, T., Gelfand, I.M., Feder, H.H., and Bonder, E.M. (2004). Rho overexpression leads to mitosis-associated detachment of cells from epithelial sheets: a link to the mechanism of cancer dissemination. Proc Natl Acad Sci U S A 101, 12526–12530.

25. Pease, J.C., and Tirnauer, J.S. (2011). Mitotic spindle misorientation in cancer--out of alignment and into the fire. J Cell Sci 124, 1007–1016. 10.1242/jcs.081406.

26. Hanahan, D., and Weinberg, R.A. (2000). The hallmarks of cancer. Cell 100, 57–70.

27. Peglion, F., and Etienne-Manneville, S. (2023). Cell polarity changes in cancer initiation and progression. Journal of Cell Biology 223.

28. Cox, T.R., and Erler, J.T. (2011). Remodeling and homeostasis of the extracellular matrix: implications for fibrotic diseases and cancer. Dis Model Mech 4, 165–178.

29. Krahn, M.P., Bückers, J., Kastrup, L., and Wodarz, A. (2010). Formation of a Bazooka– Stardust complex is essential for plasma membrane polarity in epithelia. Journal of Cell Biology 190, 751–760.

30. Humbert, P.O., Grzeschik, N.A., Brumby, A.M., Galea, R., Elsum, I., and Richardson, H.E. (2008). Control of tumourigenesis by the Scribble/Dlg/Lgl polarity module. Oncogene 27, 6888–6907.

31. Oda, H., and Takeichi, M. (2011). Structural and functional diversity of cadherin at the adherens junction. Journal of Cell Biology 193, 1137–1146.

32. Eble, J.A., and Niland, S. (2019). The extracellular matrix in tumor progression and metastasis. Clinical & experimental metastasis 36, 171–198.

33. Conlon, G.A., and Murray, G.I. (2019). Recent advances in understanding the roles of matrix metalloproteinases in tumour invasion and metastasis. The Journal of pathology 247, 629–640.

34. Murray, M.A., Fessler, L.I., and Palka, J. (1995). Changing distributions of extracellular matrix components during early wing morphogenesis in Drosophila. Developmental biology 168, 150–165.

35. Leitão, A.B., and Sucena, É. (2015). Drosophila sessile hemocyte clusters are true hematopoietic tissues that regulate larval blood cell differentiation. eLife 4, e06166.

36. Crozatier, M., and Meister, M. (2007). Drosophila haematopoiesis. Cell Microbiol 9, 1117–1126..

37. Harrison, D.A., Binari, R., Nahreini, T.S., Gilman, M., and Perrimon, N. (1995). Activation of a Drosophila Janus kinase (JAK) causes hematopoietic neoplasia and developmental defects. The EMBO Journal 14, 2857–2865-2865.

38. Ladányi, A. (2015). Prognostic and predictive significance of immune cells infiltrating cutaneous melanoma. Pigment Cell & Melanoma Research 28, 490–500.

39. Méthot, N., and Basler, K. (1999). Hedgehog controls limb development by regulating the activities of distinct transcriptional activator and repressor forms of Cubitus interruptus. Cell 96, 819–831.

40. Ng, M., Diaz-Benjumea, F.J., Vincent, J.-P., Wu, J., and Cohen, S.M. (1996). Specification of the wing by localized expression of wingless protein. Nature 381, 316–318.

41. Hanahan, D., and Weinberg, R.A. (2011). Hallmarks of cancer: the next generation. Cell 144, 646–674.

42. Bartek, J. (2011). DNA damage response, genetic instability and cancer: from mechanistic insights to personalized treatment. Molecular oncology 5, 303.

43. Yao, Y., and Dai, W. (2014). Genomic instability and cancer. Journal of carcinogenesis & mutagenesis 5, 1000165.

44. Bageritz, J., Willnow, P., Valentini, E., Leible, S., Boutros, M., and Teleman, A.A. (2019). Gene expression atlas of a developing tissue by single cell expression correlation analysis. Nature Methods 16, 750–756.

45. Deng, M., Wang, Y., Zhang, L., Yang, Y., Huang, S., Wang, J., Ge, H., Ishibashi, T., and Yan, Y. (2019). Single cell transcriptomic landscapes of pattern formation, proliferation and growth in Drosophila wing imaginal discs. Development 146.

46. Everetts, N.J., Worley, M.I., Yasutomi, R., Yosef, N., and Hariharan, I.K. (2021). Single-cell transcriptomics of the Drosophila wing disc reveals instructive epithelium- to-myoblast interactions. Elife 10, e61276.

47. Becht, E., McInnes, L., Healy, J., Dutertre, C.A., Kwok, I.W.H., Ng, L.G., Ginhoux, F., and Newell, E.W. (2019). Dimensionality reduction for visualizing single-cell data using UMAP. Nat Biotechnol 37, 38–44.

48. Gunage, R.D., Reichert, H., and VijayRaghavan, K. (2014). Identification of a new stem cell population that generates Drosophila flight muscles. Elife 3, e03126.

49. Sudarsan, V., Anant, S., Guptan, P., VijayRaghavan, K., and Skaer, H. (2001). Myoblast diversification and ectodermal signaling in Drosophila. Dev Cell 1, 829–839.

50. Klambt, C., Glazer, L., and Shilo, B.Z. (1992). breathless, a Drosophila FGF receptor homolog, is essential for migration of tracheal and specific midline glial cells. Genes Dev 6, 1668–1678.

51. Roy, S., Huang, H., Liu, S., and Kornberg, T.B. (2014). Cytoneme-mediated contact- dependent transport of the Drosophila decapentaplegic signaling protein. Science 343, 1244624.

52. Worley, M.I., Everetts, N.J., Yasutomi, R., Chang, R.J., Saretha, S., Yosef, N., and Hariharan, I.K. (2022). Ets21C sustains a pro-regenerative transcriptional program in blastema cells of Drosophila imaginal discs. Current biology 32, 3350–3364. e3356.

53. Martin-Blanco, E., Gampel, A., Ring, J., Virdee, K., Kirov, N., Tolkovsky, A.M., and Martinez-Arias, A. (1998). puckered encodes a phosphatase that mediates a feedback loop regulating JNK activity during dorsal closure in Drosophila. Genes Dev 12, 557–570.

54. Garcia-Arias, J.M., Pinal, N., Cristobal-Vargas, S., Estella, C., and Morata, G. (2023). Lack of apoptosis leads to cellular senescence and tumorigenesis in Drosophila epithelial cells. Cell Death Discovery 9, 281.

55. Pinal, N., Calleja, M., and Morata, G. (2019). Pro-apoptotic and pro-proliferation functions of the JNK pathway of Drosophila: roles in cell competition, tumorigenesis and regeneration. Open Biol 9, 180256.

56. Karkali, K., Pastor-Pareja, J.C., and Martin-Blanco, E. (2023). JNK signaling and integrins cooperate to maintain cell adhesion during epithelial fusion in Drosophila. Front Cell Dev Biol 11, 1034484.

57. Joyce, E.F., Pedersen, M., Tiong, S., White-Brown, S.K., Paul, A., Campbell, S.D., and McKim, K.S. (2011). Drosophila ATM and ATR have distinct activities in the regulation of meiotic DNA damage and repair. J Cell Biol 195, 359–367.

58. Canman, C.E., and Lim, D.-S. (1998). The role of ATM in DNA damage responses and cancer. Oncogene 17, 3301–3308.

59. Virchow, R. (1858). Die cellularpathologie in ihrer begründung auf physiologische und pathologische gewebelehre. Zwanzig vorlesungen gehalten während der monate februar, märz und aprilim Pathologischen institute zu Berlin (A. Hirschwald).

60. Alter, N.M. (1925). Mechanical irritation as etiologic factor of cancer: clinical observation. The American Journal of Pathology 1, 511.

61. Byun, J.S., and Gardner, K. (2013). Wounds that will not heal: pervasive cellular reprogramming in cancer. Am J Pathol. 182, 1055–1064.

62. Deyell, M., Garris, C.S., and Laughney, A.M. (2021). Cancer metastasis as a non- healing wound. British Journal of Cancer 124, 1491–1502.

63. Vidal, M., and Cagan, R.L. (2006). Drosophila models for cancer research. Current opinion in genetics & development 16, 10–16.

64. Fox, D.T., Cohen, E., and Smith-Bolton, R. (2020). Model systems for regeneration: Drosophila. Development 147, dev173781.

65. Hariharan, I.K., and Serras, F. (2017). Imaginal disc regeneration takes flight. Current opinion in cell biology 48, 10–16.

66. Fan, Y., and Bergmann, A. (2008). Apoptosis-induced compensatory proliferation. The Cell is dead. Long live the Cell! Trends Cell Biol 18, 467–473.

67. Ryoo, H.D., Gorenc, T., and Steller, H. (2004). Apoptotic cells can induce compensatory cell proliferation through the JNK and the Wingless signaling pathways. Dev Cell 7, 491–501.

68. Perez-Garijo, A., Martin, F.A., and Morata, G. (2004). Caspase inhibition during apoptosis causes abnormal signalling and developmental aberrations in Drosophila. Development 131, 5591–5598.

69. Goenka, A., Khan, F., Verma, B., Sinha, P., Dmello, C.C., Jogalekar, M.P., Gangadaran, P., and Ahn, B.C. (2023). Tumor microenvironment signaling and therapeutics in cancer progression. Cancer Communications 43, 525–561.

70. Gerlach, S.U., and Herranz, H. (2020). Genomic instability and cancer: lessons from Drosophila. Open Biol 10, 200060.

71. Fujiwara, T., Bandi, M., Nitta, M., Ivanova, E.V., Bronson, R.T., and Pellman, D. (2005). Cytokinesis failure generating tetraploids promotes tumorigenesis in p53-null cells. Nature 437, 1043–1047.

72. Ciccia, A., and Elledge, S.J. (2010). The DNA damage response: making it safe to play with knives. Mol Cell 40, 179–204.

73. Srivastava, N., Gochhait, S., de Boer, P., and Bamezai, R.N.K. (2009). Role of H2AX in DNA damage response and human cancers. Mutat Res 681, 180–188. 1

74. Bi, X., Gong, M., Srikanta, D., and Rong, Y.S. (2005). Drosophila ATM and Mre11 are essential for the G2/M checkpoint induced by low-dose irradiation. Genetics 171, 845–847.

75. Brodsky, M.H., Nordstrom, W., Tsang, G., Kwan, E., Rubin, G.M., and Abrams, J.M. (2000). Drosophila p53 binds a damage response element at the reaper locus. Cell 101, 103–113.

76. Peters, M., DeLuca, C., Hirao, A., Stambolic, V., Potter, J., Zhou, L., Liepa, J., Snow, B., Arya, S., Wong, J., et al. (2002). Chk2 regulates irradiation-induced, p53-mediated apoptosis in Drosophila. Proc Natl Acad Sci U S A 99, 11305–11310.

77. Martin, F.A., Perez-Garijo, A., and Morata, G. (2009). Apoptosis in Drosophila: compensatory proliferation and undead cells. Int J Dev Biol 53, 1341–1347.

78. Martin-Belmonte, F., and Perez-Moreno, M. (2012). Epithelial cell polarity, stem cells and cancer. Nature Reviews Cancer 12, 23–38.

79. La Marca, J.E., and Richardson, H.E. (2020). Two-faced: roles of JNK signalling during tumourigenesis in the Drosophila model. Frontiers in cell and developmental biology 8, 42.

80. Misra, J.R., and Irvine, K.D. (2018). The Hippo Signaling Network and Its Biological Functions. Annu Rev Genet 52, 65–87.

81. Sun, G., and Irvine, K.D. (2011). Regulation of Hippo signaling by Jun kinase signaling during compensatory cell proliferation and regeneration, and in neoplastic tumors. Dev Biol 350, 139–151.

82. Doggett, K., Grusche, F.A., Richardson, H.E., and Brumby, A.M. (2011). Loss of the Drosophila cell polarity regulator Scribbled promotes epithelial tissue overgrowth and cooperation with oncogenic Ras-Raf through impaired Hippo pathway signaling. BMC Dev Biol 11, 57.

83. Gerlach, S.U., Eichenlaub, T., and Herranz, H. (2018). Yorkie and JNK Control Tumorigenesis in Drosophila Cells with Cytokinesis Failure. Cell Rep 23, 1491–1503.

84. Casas-Tinto, S., Lolo, F.N., and Moreno, E. (2015). Active JNK-dependent secretion of Drosophila Tyrosyl-tRNA synthetase by loser cells recruits haemocytes during cell competition. Nat Commun 6, 10022.

85. Motoyama, N., and Naka, K. (2004). DNA damage tumor suppressor genes and genomic instability. Current opinion in genetics & development 14, 11–16.

86. Liu, Y., Dantas, E., Ferrer, M., Miao, T., Qadiri, M., Liu, Y., Comjean, A., Davidson, E.E., Perrier, T., Ahmed, T., et al. (2025). Hepatic gluconeogenesis and PDK3 upregulation drive cancer cachexia in flies and mice. Nat Metab 7, 823–841.

